# Social navigation: distance and grid-like codes support navigation of abstract social space in human brain

**DOI:** 10.1101/2023.05.12.538784

**Authors:** Zilu Liang, Simeng Wu, Jie Wu, Wenxu Wang, Shaozheng Qin, Chao Liu

## Abstract

People form impressions about others during daily social encounters and infer personality traits from others’ behaviors. Such trait inference is thought to rely on two universal dimensions, i.e., competence and warmth. These two dimensions can be used to construct a ‘social cognitive map’ organizing massive information obtained from social encounters efficiently. Originated from spatial cognition, the neural codes supporting representation and navigation of spatial cognitive map has been widely studied. Recent studies suggest similar neural mechanism subserves the map-like architecture in social cognition as well. Here we investigated how spatial codes operate beyond physical environment and support the representation and navigation of social cognitive map. We designed a social value space defined by two dimensions of competence and warmth. Behaviorally, participants were able to navigate to a learned location from random starting locations in this abstract social space. At neural level, we identified representation of distance in precuneus, fusiform gyrus and middle occipital gyrus. We also found partial evidence of grid-like representation patterns in medial prefrontal cortex and entorhinal cortex. Moreover, the intensity of grid-like response scaled with performance of navigating in social space and social avoidance trait scores. Our findings suggest a neurocognitive mechanism by which social information can be organized into a structured representation namely cognitive map and its relevance to social well-being.

## Introduction

Countless daily social encounters pose people with the need to organize information about social encounters efficiently. Though people may ascribe a variety of traits to others, these inferences has been identified to rely on a few universal dimensions (Fiske et al., 2007; Stolier et al., 2020). The most widely recognized dimensions are warmth, which concerns others intentions, and competence, which concerns others capability and possession of resources (Cuddy et al., 2008). These universal dimensions can be used to construct a structured representation organizing impressions of other people or other social information alike, just like how one would specify a location in a plane using orthogonal dimensions for spatial navigation. Analogous to Tolman’s original concept (Tolman, 1948), here we termed this structured representation of social information about other people as ‘social cognitive map’, which may form the basis for social cognition. Having such a cognitive map for social perception is crucial. It helps us organize various experiences, track updates and guide novel inferences efficiently (Son et al., 2021). Despite its importance in navigating social world, its neural underpinning remains under-investigated.

The resemblance between social and spatial cognitive map has led to the proposition that social cognitive map is also supported by similar mechanisms that map physical space (Schafer & Schiller, 2018). In the domain of spatial navigation, it has been demonstrated that entorhinal grid cell activity supports representational coding of cognitive map. Direct recordings in rodents (Hafting et al., 2005) and humans (Jacobs et al., 2013) have found that the firing locations of grid cells form periodic triangular that covers the entire available arena during spatial navigation task. In this sense, grid cells form the basic units of a neural map of the spatial environment. One of the prominent properties of grid cells is their consistent grid orientation (i.e., the orientation of the grid relative to the environment) shared by neighboring grid cells (Hafting et al., 2005; Sargolini et al., 2006). Moreover, the overlapping population formed by grid cells and head direction cells as well as conjunctive grid × head-direction cells in deeper layers of entorhinal cortex enables directional as well as positional tuning of the firing rate (Sargolini et al., 2006). It has been proposed that this conjunctive representation at a population level can yield a directionally modulated firing pattern with hexagonal periodicity in which population activity is enhanced when agent’s moving direction is aligned to the grid axes (Kriegeskorte & Storrs, 2016; Kunz et al., 2019). Human studies using non-invasive neuroimaging techniques rely on this directional preference of populational conjunctive grid × head-direction cell activity to detect evidence of grid-like code in fMRI BOLD signals during spatial navigation (Doeller et al., 2010). Follow-up studies have reported entorhinal grid code during virtual as well as imagined navigation (Horner et al., 2016), and even mental simulation (Bellmund et al., 2016). More importantly, pioneering studies showed that the function of grid-like code goes beyond the scope of spatial navigation. Grid-like codes in entorhinal cortex and prefrontal cortex have been observed during nonspatial navigation such as perceptual (Aronov et al., 2017; Bao et al., 2019; Killian et al., 2012; Nau et al., 2018), conceptual (Constantinescu et al., 2016), semantic space (Vigano & Piazza, 2020; Vigano et al., 2021) and more recently, discrete social hierarchy (Park et al., 2021). Evidence from above studies converges into a domain-general role of grid-like codes in organizing cognitive maps of various types including spatial and nonspatial domains (Bellmund et al., 2018).

In this study, we tested the above hypothesis that social cognitive map is supported by a grid cell-like coding mechanism observed in spatial navigation. First, leveraging the theoretical framework of the stereotype content model, we set up an abstract two-dimensional social value map which are composed of competence and trustworthiness dimensions of social perception. Next, we adapted paradigms used in human fMRI grid-code study to test the hypothesis that grid-like codes and distance codes resembling those underpinning spatial navigation are involved in navigation in this abstract social space. Finally, to complete the argument that neural codes for social navigation subserve social cognition, we explored the link between neural codes during social navigation and social skills measured by navigational performance and social trait scores.

## Results

To look for neural underpinnings of navigation in an abstract social space, we adapted a set of tasks from previous studies illustrating the relevance of grid-like code in navigating abstract concept space (Constantinescu et al., 2016). Participants received intensive training on navigating in this abstract social space with precision. They completed a learning session, a review session during behavioral training on the first day, and a scanning session on the second day.

We designed the social value map, an abstract social space structuring one’s social perception of other people’s social value. It was defined by two ecologically important dimensions: competence and trustworthiness. A visualization with two adjacent bars enclosed by a square box was designed to represent the location on this social value map, with the height of each bar representing value of each dimension. We operationally defined these two dimensions under the framework of an investment game, in which trustee’s social value is determined by the ability to earn profit (i.e., profit rate as competence) and the proportion of earnings returned to the investor (i.e., return rate as trustworthiness). The range of competence and trustworthiness was limited to between 0 and 1 to make sure that they are realistic (i.e., it is unlikely that trustees will return you more than they have gained), and more importantly, orthogonal and share same metric system. Participants played the role of an investor in this investment task (**Figure S1A, Supplementary Methods 1.2**) at the very beginning of the experiment to develop a concrete understanding of the visual analogue and the quantitative meaning of these two dimensions.

Six avatars with different levels of competence and trustworthiness **(Figure 1A)**, analogous to landmarks in the physical environment, were placed on the social value map of size 1 unit × 1 unit in accordance with the range of competence and trustworthiness. The current spatial arrangement aimed to make avatars distinguishable on both dimensions while spreading widely across the whole space. Specifically, each participant’s set of avatars’ coordinates was sampled within circles of 1/30-unit radius around center coordinates (**Figure 1B**). In the experiment, avatars were represented by passport-style photos of faces of volunteers taken against a plain blue background. Photographs were matched on competence and trustworthiness ratings based on data from an independent online sample **(Supplementary Methods 1.1**). Additionally, scanned participants completed ratings on the selected faces before and after the experiment to reassure that there was no pre-existing bias in social perception of the avatars and to test if they updated social perception according to learned characteristics of the avatars **(Supplementary Methods 1.9**).

**Figure 1.**
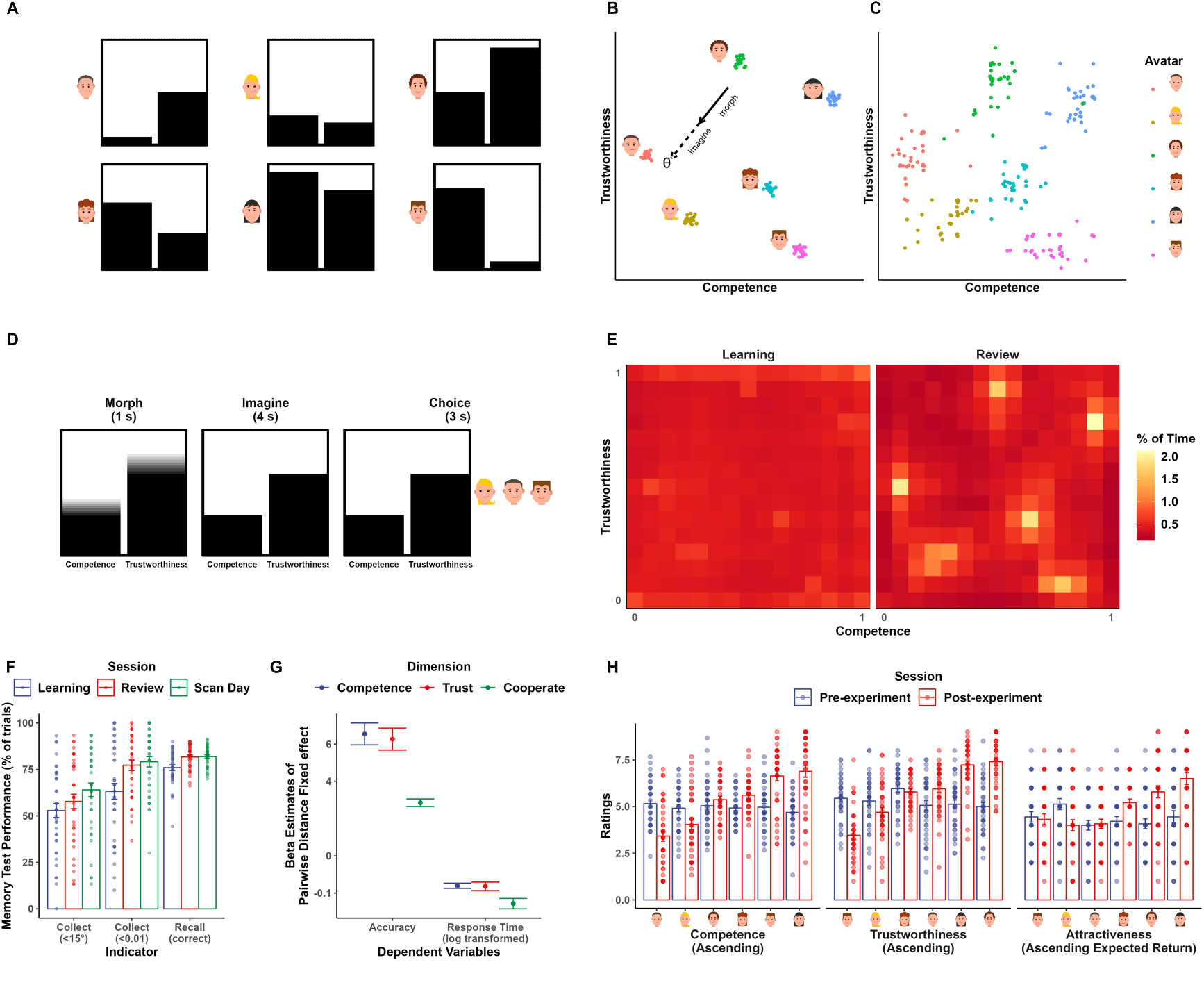
Experimental Design and Behavioral Performance. (**A**) Visual analogues illustrating each avatar’s social values. Height of the left bars signals values on competence dimension, while height of the right bars signals trustworthiness. Labels for each of the two bars are omitted for illustration purpose. (**B**) Corresponding layout of the social value map and example trajectory in recall task. Scattered dots indicate the actual sets of positions generated for participants. (**C**) Positions of the avatars indicated by participants at the end of the whole experiment. (**D**) Trial timeline of the recall task. Participants first watched the bars morphing for one second according to a predefined competence-trustworthiness ratio (i.e., the first half of a trajectory in Figure 1B). After the bars stopped morphing, they were instructed to imagine the bars continue to morph according to the same ratio, at the same speed and for the same amount of time. (i.e., the second half of a trajectory in Figure 1B) (**E**) Color-coded trajectory map of the explore task during learning and review session. Color indicated the percentage of time spent in each pixel of the social value map during the explore task in the learning (left panel) and review (right panel) session. A pattern emerged that participants spent more time at avatars and less time at edge during review. (**F**) Memory test performance of the collect and recall task in learning, review and scan-day session. Collect(<15°): percentage of trials where deviation of first transition from ideal trajectory was smaller than 15°; Collect(<0.01): percentage of trials where distance between response location and correct location was smaller than 0.01 units; Recall(correct): percentage of trials where response was correct (**G**) The distance effect in comparison task across the review and scan-day session. Colored dots indicated the estimates of distance effect regressor while error bars indicated standard error of the estimates. (**H**) Ratings became predictable from the avatars’ social value after experiment. Icons of avatars are for illustration and retrieved from https://pixabay.com/vectors/avatar-flat-modern-minimal-5261900/, https://pixabay.com/vectors/avatar-flat-modern-minimal-5261896/.

In the training, participants explored the social space by morphing the bars with different competence: trustworthiness ratio using the non-spatial controller to look for the six avatars (**Figure S1C, Supplementary Methods 1.4**). Morphing the bars resembles making a transition in the social space in that it was done by two steps: 1) deciding a trajectory to follow by changing the competence: trustworthiness ratio using the non-spatial controller, 2) how further along to follow that trajectory. No prior information about the characteristics of the avatars was provided to the participants. Thus, participants could only explore social space as if they were looking for landmarks in a newly introduced physical environment. An avatar would pop out when the visualization matched his/her characteristics. Specifically, the avatar would pop out when participants’ current location fell within a 0.01-unit radius of the correct location on the social value map. In this way, participants not only learned the characteristics of each avatar but also became familiar with the whole space even though this map-like structure was never revealed to them.

After participants were fully acquainted with the avatars and the social space, a set of memory tasks were used to test if participants learned the avatars, and more importantly, if they formed an internal representation of the social value map, even though the map-like structure was never directly revealed to them. Participants never received feedback on these memory test tasks, so they had to rely on knowledge learned in the explore task. The first one was the collect task (**Figure S1D, Supplementary Methods 1.5**) which required participant to navigate to the location of the avatars from random starting locations in the abstract space. In each trial, they were instructed to morph the box on the left to match the characteristic of the avatar shown on the right. Specifically, we asked participants to make as few transitions as possible. Two performance indices were calculated: 1) the deviation of participant’s first transition from the ideal trajectory; and 2) the Euclidean distance between the location corresponding to participants’ morphed bars and the avatar’s actual location in the abstract space (**Supplementary Methods 2.1.2**). The second one was the recall task, which required participant to mentally traverse the abstract social space according to a half-seen trajectory and identify the destination that follows (**Figure 1D, Supplementary Methods 1.6**). In each trial, participants were first shown the two bars morphing according to a predefined competence: trustworthiness ratio for 1 second. The bars then stopped morphing, and participants were instructed to imagine the bars keep morphing according to the same ratio, at the same speed, and for the same amount of time. After this, participants had to choose which of the three given options matched the bars after imagination. Essentially, each trial is a trajectory vector defined by moving direction (i.e., morphing ratio) and travelled distance on the social value map. We specifically sample the trajectories from a uniform distribution (**Figure S8**). We investigated the neural representation of the trajectories while participant performed this task. The third one was the compare task (**Figure S1E, Supplementary Methods 1.8**) which required participants to compare different pairs of avatars on one given dimension (competence and trustworthiness) or by a fair combination of both dimensions (willingness to cooperate). Assuming distance is a measure of similarity between two avatars, the closer two avatars are, the more similar they are, hence distinguishing them will be harder, which results in less accurate response and longer reaction time (similar to the inferential distance effect in ordered relationships (Potts, 1974)). If participants had map-like representations, then their response in this task would show such correlation pattern with distance in the 2D social space .The final one was a map task at the end of the experiment in which participants were informed about the map-like structure and were asked to indicate the location of each avatar on an empty social value map using mouse click.

### Participants construct social value map after associative learning of avatars and corresponding characteristics

Participants were successful in reconstructing the designed layout if avatars in a map task at the end of experiment (**Figure 1C**). Linear mixed effect models revealed that participants’ performance in tasks significantly improved over sessions. In the explore task, it took them significantly less time to find all avatars in the review session compared to the learning session (β_session_ = -67.589, t = -37.604, p < 0.001). They also spent much less time at edge (β = -0.104, t = -9.070, p < 0.001) and more time at avatars (β_session_ = 0.023, t = 7.367, p < 0.001) during exploration in the review session compared to the learning session (**Figure 1E, Supplementary Methods 2.1.1**). Likewise, there was evident improvement in collect and recall task as training progressed (**Figure 1F**, Percentage of first transition deviation < 15° in collect task: β_session_ = 0.050411, t = 3.2187, p = 0.002; Percentage of distance < 0.01 in collect task: β_session_ = 0.047, t = 3.002, p = 0.003; Accuracy in recall task: β_session_ = 0.020, t = 4.213, p < 0.001). Furthermore, participants were able to form an implicit map-like structure and integrate both dimensions when making decisions between avatars. In the compare task, linear mixed effect models (**Supplementary Methods 2.1.4**) revealed significant effect of distance between avatars in the compared dimension on response time and accuracy as predicted (**Figure 2G**, all p<0.001). Lastly, learning induced changes in participants’ ratings on avatars on all rating items (competence, trustworthiness, and attractiveness) (**Figure 2H, Table S1A**). There was no preference over the six presented faces when participants were naïve to the social value of different avatars (**Table S1B**, all p>0.890), but their ratings became significantly dissociable according to the learned social value at the end of the experiment (**Table S1B**, all p<0.001). These results indicated successful formation of a 2D social map by associating each avatar with respective social value.

**Figure 2.**
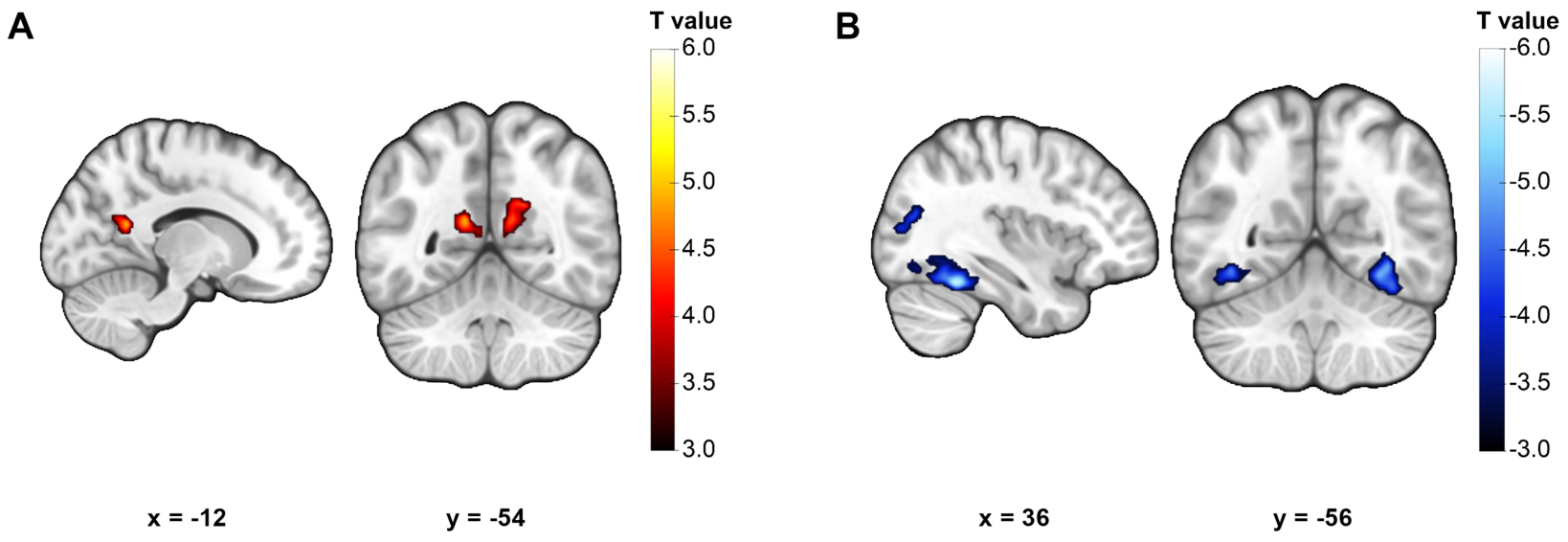
Neural representation of Euclidian distance on the social value map. (**A**) Activity in the bilateral Precuneus positively correlated with travelled Euclidian distance. (**B**) Activity in bilateral Fusiform and the right middle occipital gyrus negatively correlated with travelled Euclidian distance. Display threshold: cluster-defining threshold p < 0.001.

### Precuneus and fusiform jointly encode Euclidian distance during social navigation

Next, we investigated the neural representations supporting the social cognitive map. To examine the neural code for distance information, we conducted whole-brain analysis with a GLM with two regressors modelling the morph stage and choice stage. Travelled Euclidian distance (i.e., the length of each trial’s trajectory vector) was entered as parametric modulator of the morph stage regressor (**Supplementary Methods 2.2.3**). Even though we did not specifically control for the distribution of travelled distance, there was quite a lot trial-wise variability that allowed us to conduct this test (**Figure S9**).

This analysis revealed two marginally significant clusters in the precuneus whose activity positively correlated with travelled distance on the social value map (**Figure 2A, Table S2A**, right precuneus: peak coordinate [x, y, z] = [12, -56, 26], t(37) = 4.952, p = 0.056 FWE-corrected; left precuneus: peak coordinate [x, y, z] = [-12, -54, 18], t(37) = 4.530, p = 0.058 FWE-corrected; cluster-defining threshold p < 0.001 FWE-corrected). Interestingly, we also observed negative correlation between travelled distance and activity of clusters in several regions, including bilateral fusiform gyrus (**Figure 2B, Table S2B**, right fusiform: peak coordinate [x, y, z] = [36, -48, -20], t(37) = 5.912, p < 0.001 FWE-corrected; left fusiform: peak coordinate [x, y, z] = [-38, -56, -10], t(37) = 4.724, p = 0.008 FWE-corrected) and the right middle occipital gyrus (**Figure 2B, Table S2B**, peak coordinate [x, y, z] = [34, -68, 20], t(37) = 4.649, p = 0.018 FWE-corrected; cluster-defining threshold p < 0.001).

### Grid-like activity aligned to prefrontal and entorhinal grid orientation

To look for grid-like activity, we implemented the orientation-estimation approach (Bellmund et al., 2018; Doeller et al., 2010) based on univariate analysis. First, a whole-brain quadrature filter analysis was conducted which served as a functional localizer to identify regions sensitive to hexagonal modulation, independent of grid orientation (**Supplementary Methods 2.2.1**). This GLM had two regressors modelling the morph stage and choice stage. For the morph stage regressor, we included two parametric modulators corresponding to the sixfold sinusoidal six-fold modulation of trajectory direction (i.e., sin6θ and cos6θ, θ is the direction). Next, we defined spherical ROIs with a 5-mm radius centered at the peak coordinate of each significant cluster (**Table S3**) to estimate the grid orientation of each ROI. As no region in the entorhinal cortex survived this analysis, we additionally defined four entorhinal ROI based on subdivisions from anatomical mask (Maass et al., 2015). Finally, we conducted the leave-one-out cross-validation analysis to test for hexagonal modulation aligned to grid orientation of the ROIs for each participant, i.e., the grid orientation consistency effect (**Supplementary Methods 2.2.2**). In this cross-validation approach, we estimated grid orientation using data from three runs and tested the grid orientation consistency effect in the held-out run. Grid consistency effect was tested using a GLM that classified trials into 12 bins according to its trajectory direction offset from the estimated grid orientation, and built a contrast to test if activity is stronger in aligned trials than in misaligned trials (**Figure 3A)**.

**Figure 3.**
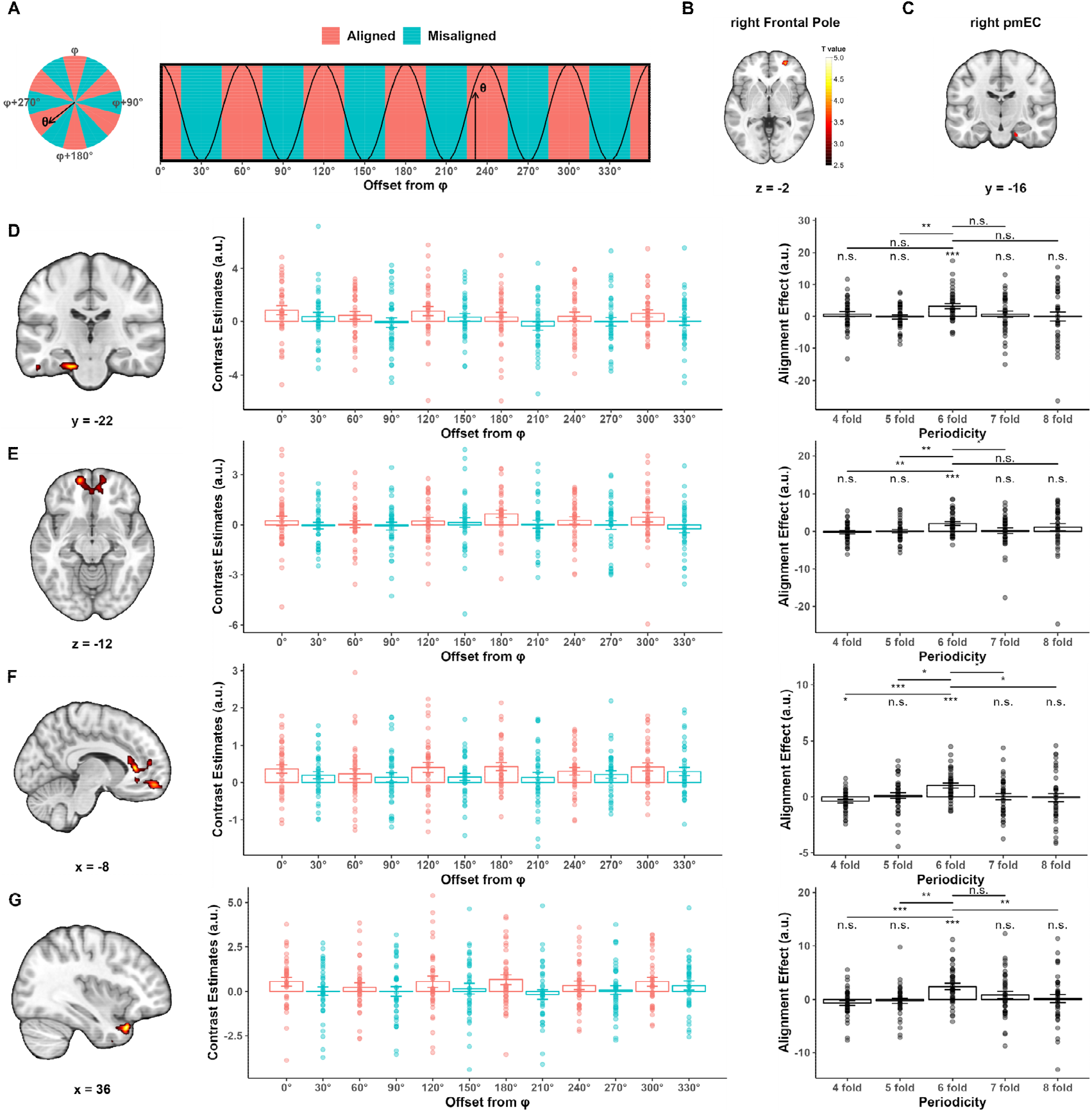
Evidence of grid-like activity aligned to putative grid orientation in the right frontal pole and the right posterior-medial entorhinal cortex. **(A)** Theoretical prediction of grid-like activity **(B-C)** ROIs for deriving putative grid orientations: **(B)** right FP ROI from quadrature filter analysis showing sensitivity to hexagonal modulation. A 5-mm sphere was defined around the peak coordinate to compute grid angle. Display threshold: voxel-level p < 0.001, cluster-level p < 0.05 FWE-corrected. **(C)** Anatomically defined right pmEC ROI used to compute grid angle. **(D-G)** Grid like activity aligned to putative grid orientations in the right FP ROI (**D-E**) and right pmEC (**F-G**) ROI respectively. Left panels: clusters from whole-brain hexagonal consistency analysis. Color indicates T statistics as shown in the colorbar in **B**. Display threshold: voxel-level p < 0.005, cluster-level p < 0.05 FWE-corrected. Middle panels: hexagonal consistency effects plotted as contrast estimates of the 12 trial-bin regressors extracted from corresponding cluster in the left panel; To illustrate effect in EC in E, estimates were extracted from the intersection of the suprathreshold cluster and anatomical mask of EC. Right panels: such effects were specific to six-fold. Notes: n.s., p>0.05, *p<0.05, **p<0.01, ***p<0.001. Abbreviations: FP, frontal pole; pmEC, posterior-medial entorhinal cortex; vmPFC, ventral medial prefrontal cortex; STP, superior temporal pole.

Of all the grid orientations derived from the above ROIs, we only found grid consistency effect aligned to the grid orientation of the ROI in the right frontal pole (right FP) cluster (**Figure 3B**, peak coordinate [x, y, z] = [26, 52, -2], t(37) = 4.494, cluster-level p = 0.017 FWE-corrected, cluster-defining threshold p < 0.001) and the grid orientation of the anatomically defined right posterior-medial entorhinal cortex (right pmEC, **Figure 3C**). For the right FP grid orientation, whole-brain analysis revealed consistency effect in bilateral ventral medial prefrontal cortex (vmPFC, **Figure 3D** left and middle panel, peak coordinate [x, y, z] = [-14, 56, -12], t(37) = 4.245, cluster-level p = 0.033 FWE-corrected, cluster-defining threshold p < 0.005) and the left entorhinal cortex (left EC) extending to the parahippocampus (**Figure 3E** left and middle panel, peak coordinate [x, y, z] = [-16, -22, -26], t(37) = 4.563, cluster-level p = 0.024 FWE-corrected, cluster-defining threshold p < 0.005). For the right pmEC grid orientation, significant consistency effect was found in two clusters in bilateral vmPFC (**Figure 3F** left and middle panel, right rectus: peak coordinate [x, y, z] = [10, 34, -18], t(37) = 4.908, cluster-level p = 0.033 FWE-corrected; left ACC: peak coordinate [x, y, z] = [-8, 32, 6], t(37) = 4.810, cluster-level p = 0.009 FWE-corrected; cluster-defining threshold p < 0.005) and one cluster in right superior temporal pole (right STP, **Figure 3G** left and middle panel, peak coordinate [x, y, z] = [36, 20, -30], t(37) = 4.962, cluster-level p = 0.018 FWE-corrected, cluster-defining threshold p < 0.005). However, we did not find evidence of grid-like code in the entorhinal cortex aligned to its own putative grid orientation either with this orientation estimation approach (**Figure S5A-B**) or with the representation similarity analysis approach (**Figure S5C-D, Supplementary Methods 2.3**).

One important assumption underlying the univariate analysis for grid-like code in human fMRI data is neighboring grid cells share similar grid orientation. Consistent with previous studies, we examined this assumption in our dataset by testing the distribution of putative voxel-wise grid orientations in the right FP ROI and in the right pmEC ROI for each participant (**Supplementary Methods 2.4**). This revealed a clustered distribution of voxel-wise grid orientations in FP ROI (V-test, all p < 0.011, except in one participant’s one estimating set p = 0.176; see **Figure S2A** for representative participants). Previous studies have also shown that grid orientations across participants tend to distribute uniformly. This has been replicated in the right FP ROI in our dataset (**Figure S3** left panel, Rayleigh’s tests for nonuniformity, all p > 0.782). However, in the pmEC ROI, these results are less consistent. With regards to voxel-wise orientations, 31 participants show the above clustered distribution (V-test, all p < 0.040; see **Figure S2B** for representative participants), but there are 7 participants each of whom has at least one estimating set that didn’t pass the statistical test (V-test, all p > 0.068). Across participants, only grid orientations from two estimating sets conformed to the uniform distribution (**Figure S3** right panel, Rayleigh’s tests for nonuniformity, run 1,2,3: p = 0.010, run 1,2,4: p =0.019).

For completeness, we tested the specificity of hexagonal modulation. That is, the identified alignment effect only exists with six-fold periodicity. The above orientation estimation and cross-validation procedure was repeated for each of the four controlled periodicity (i.e., 4-fold, 5-fold, 7-fold, and 8-fold). Signals were extracted from the clusters that showed significant hexagonal consistency. One-sample t-tests showed that signals within these clusters were not modulated by any of the controlled periodicities and paired t-test showed that six-fold periodicity in general showed greater alignment effect (**Figure 3D-G**, right panels). In sum, our univariate analysis revealed that the vmPFC and left EC showed six-fold specific grid-like code aligned to right FP’s grid orientation and that the vmPFC and right STP showed six-fold specific grid-like code aligned to right pmEC’s grid orientation.

### Behavioral relevance of spatial codes for the social value map

Next, we want to address whether the identified distance or grid-like code could have any behavioral relevance (**Supplementary Methods 2.2.4**). Previous studies have found that hexagonal modulation effect and hexagonal consistency effect positively correlate with accuracy in the recall task in scanner (Constantinescu et al., 2016). However, we did not find such correlation in our data (**Figure S7**). Then, we explored if there is any correlation between neural indices of map-like representation and behavior performance outside the scanner as well as individual differences. The covariates we explored can be classified into two categories. The first category reflects participants ability to make decision based on the social value map, i.e., the distance effect in the compare task. Specifically, we focused on the distance effect during the cooperation block when participants choose the avatar they are more willing to cooperate with by taking into account both dimensions. This is essentially the beta weight of the distance regressor in predicting participant’s accuracy and log-transformed response time. The second category reflects participants’ social trait, including social anxiety and social avoidance scores. The covariates were entered into the second-level analysis of the consistency effect GLM separately.

Whole-brain analysis revealed significant clusters in the temporal lobe whose consistency effect scaled with distance effect in response time. For grid orientation of right FP ROI (**Figure 4A**), hexagonal consistency effect in the left middle temporal gyrus and right Lingual gyrus correlated with the distance effect (left middle temporal gyrus: peak coordinate [x, y, z] = [-36, -70, 8], t(37) = 4.528, cluster-level p < 0.001 FWE-corrected; right Lingual gyrus: peak coordinate [x, y, z] = [-38,-66, 4], t(37) = 4.369, cluster-level p < 0.001 FWE-corrected; cluster-defining threshold p < 0.005). For grid orientation of right pmEC ROI (**Figure 4B**), hexagonal consistency effect in bilateral Lingual correlated with the distance effect (left Lingual gyrus: peak coordinate [x, y, z] = [-12,-68, -2], t(37) = 4.355, cluster-level p = 0.077 FWE-corrected; right Lingual gyrus: peak coordinate [x, y, z] = [16,-62, -2], t(37) = 3.934, cluster-level p = 0.032 FWE-corrected; cluster-defining threshold p < 0.005). In general, greater distance effect in the comparison task correlated with greater hexagonal consistency in temporal lobe aligned to both prefrontal and entorhinal grid orientations. In addition, this analysis also revealed a cluster in the left precuneus/posterior cingulate cortex (left PCC, **Figure 4C**, peak coordinate [x, y, z] = [0, -64, 28], t(37) = 4.965, cluster-level p < 0.001 FWE-corrected, cluster-defining threshold p < 0.001). Its consistency effect aligned to the right FP grid orientation was negatively correlated with social avoidance score.

**Figure 4.**
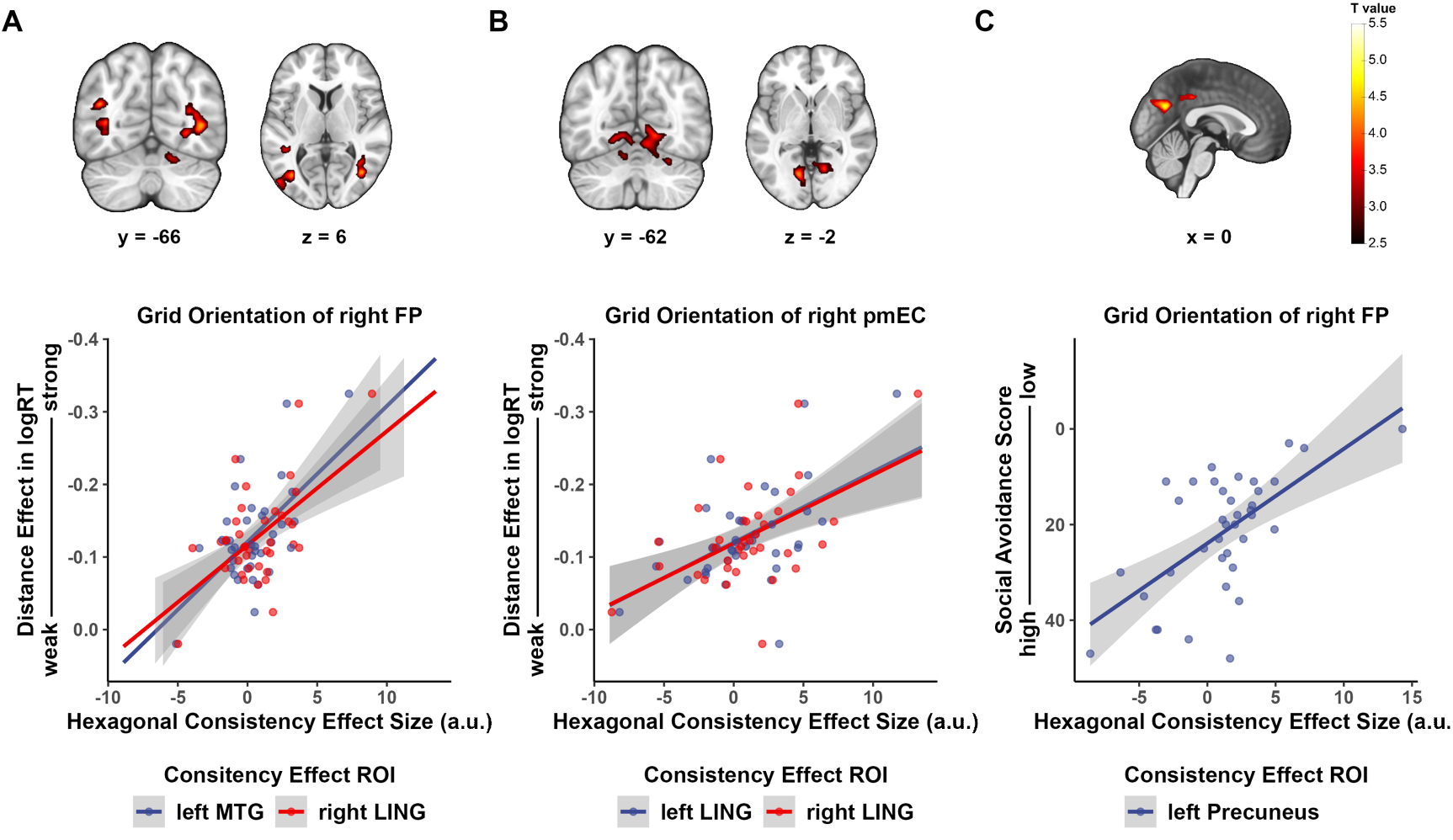
Behavioral relevance of hexagonal consistency effect. **(A-B)** Higher hexagonal consistency in temporal lobe aligned to grid orientation of **(A)** right FP ROI and **(B)** right pmEC significantly correlated with stronger distance effect in compare task when choosing preferred partners for cooperation. Display threshold: voxel-level p < 0.005, cluster-level p < 0.05 FWE-corrected. **(C)** Hexagonal consistency effect in left Precuneus aligned to grid orientation of right FP ROI significantly correlated with social avoidance score. Display threshold: voxel-level p < 0.001, cluster-level p < 0.05 FWE-corrected. Notes: n.s., p>0.05, *p<0.05, **p<0.01, ***p<0.001. Abbreviations: FP, frontal pole; pmEC, posterior-medial entorhinal cortex; MTG, middle temporal gyrus; LING, lingual gyrus.

## Discussion

In this study, we investigated whether cognitive map in social cognition domain recruit the neural processing mechanism similar to its spatial counterpart. We identified distance representation in precuneus, fusiform gyrus and middle occipital gyrus. In addition, we demonstrated some evidence of grid-like activity in prefrontal cortex and entorhinal cortex. Furthermore, we explored the behavioral relevance of grid-like activity in temporal pole and precuneus.

When participants were mentally traversing a predefined social value map, we found that activities in the bilateral precuneus were positively correlated with travelled distance, while activities in bilateral FFG and right MOG were negatively correlated with travelled distance. In spatial navigation literature, though intracranial electroencephalography studies have shown the relevance of hippocampal theta oscillation in encoding travelled distance, neural correlates of travelled distance at macroscopic level using fMRI are relatively sparsely investigated (Kunz et al., 2019). Patai et al. (2019) showed that retrosplenial cortex positively correlated with path distance in that greater distance is associated with increased precuneus activity and that this effect remained robust after controlling for Euclidean distance. Likewise, we showed that precuneus activity was positively correlated with travelled Euclidian distance during implicit navigation in the abstract social space. Moreover, the present peak coordinates coincide with those reported in a previous study by Park et al. (2020). In their study, participants learned 2D social hierarchy (competence and popularity) of two groups of agents and were asked to perform a transitive inference task in which compare a pair of agents on their competence or popularity. The authors found that bilateral precuneus represented pairwise difference in the task-relevant dimension instead of pairwise Euclidian distance in the two-dimension space. Another study reported that posterior cingulate cortex/precuneus tracks the length of the vector of social relationship change, i.e., an egocentric distance (Tavares et al., 2015). Taken together, we speculate that precuneus may encode the distance of the route, either on a concrete or an abstract map, that participants trespass under task demand. Less clear is the negative correlation we observed between travelled distance and activity in fusiform as well as middle occipital gyrus. Both FFG and MOG are involved during human spatial navigation (Boccia et al., 2014). In particular, fusiform gyrus extending into the parahippocampal place area has been showed to involve in processing spatial information such as landmark and orientation cues (Qiu et al., 2019). However, to our knowledge, there is no study that reports such negative distance coding in these occipitotemporal regions during navigation.

Apart from the distance code, another focus of the current study was to test whether grid-like codes support the organization of social knowledge. Previous studies on reporting grid-like codes mostly report hexagonal modulated pattern in the entorhinal cortex and medial prefrontal regions aligned to putative grid orientation from both regions (Constantinescu et al., 2016). Our current study partly replicated these findings, most of which are in the prefrontal region. In terms of prefrontal orientation, whole-brain analysis revealed alignment effects in the bilateral vmPFC and left EC. In terms of entorhinal orientation, however, alignment effect was found in the bilateral vmPFC and right superior temporal pole, but not in entorhinal cortex. Though we found evidence of prefrontal grid-like codes subserving social navigation, its origin remains elusive. Up to date, there has been no significant evidence of grid-like neuronal tuning in this area (Wikenheiser et al., 2021). Jacobs et al. (2013) examined grid-like cells in a variety of brain regions including the frontal cortex in neurosurgical patients during a virtual navigation task, but the proportion of significant grid-like cells in the frontal cortex did not exceed type I error rate (they did identify grid-like cells in cingulate cortex which is posterior to the prefrontal regions). Therefore, it is less likely that prefrontal grid-like codes reported in our study and previous literature come from grid cells in the prefrontal cortex. A more plausible explanation is that the anatomically and functionally connected medial- and orbitofrontal cortex and entorhinal cortex (Navarro Schroder et al., 2015; Peng et al., 2018; Squire & Zola, 1996) coordinated together to support flexible decision making. However, adopting both univariate and multivariate approaches, our current study did not find reliable effect of hexagonal grid-like code in the entorhinal region aligned to its own grid orientation, which should be evident if such hypothesis stands. Previous studies postulated that lack of univariate evidence of grid-like code may have to do with low signal-to-noise ratio (tSNR) in data from the entorhinal cortex (Bao et al., 2019). Indeed, analysis (**Supplementary Methods 2.5)** revealed lower tSNR in all four entorhinal sub regions compared to the frontal pole cluster from the quadrature filter analysis **(Figure S6A**). But we did not identify a significant relationship between tSNR and the z-transformed F statistics of hexagonal modulation either across participants or within participants (**Figure S6B-C, Table S4**). Thus, we cannot conclude that low signal-to-noise ratio in entorhinal cortex led to this null finding in our study.

Nonetheless, our study did provide some incomplete evidence of grid-like activity in the entorhinal cortex and prefrontal region while participants were mentally trespassing an abstract social space. Grid codes are proposed to provide a metric of space (Moser & Moser, 2008) and underly the process of path integration (McNaughton et al., 2006). Previous studies have suggested that entorhinal grid codes are capable of representing space even at a higher level of abstraction that goes beyond spatial navigation to support cognitive flexibility in general (Bellmund et al., 2018). Existing literature revealed correlation between entorhinal and prefrontal grid-like code and performance in spatial, perceptual and conceptual space. Indeed, our exploratory analysis showed that the univariate hexagonal consistency effect in temporal lobe scaled with the distance effect in a comparison task block when participants chose their preferred partner for future cooperation. The distance effect reflected the extent to which participants relied their preference judgment on the pairwise distance between given pair of avatars. As we explicitly required participants to take both dimension into consideration, it reflected participant’s ability to plan a route between landmarks (which represents retained impression of social encounters) in the abstract social space. This is very similar to what Park et al. (2021) reported in their recent study. In this study, participants were trained on the rankings of 16 individuals on two dimensions (competence and popularity) separately, then they were asked to make comparisons between novel pairings unseen during training. Replicating their previous findings (Park et al., 2020), the authors found that the entorhinal and hippocampus system successfully reconstruct the unseen whole spectrum of social hierarchies as a 2D cognitive map. Moreover, they found that grid-like representations support trajectories of novel inferences on this 2D cognitive map that underpins decision making. Park’s study and ours differed in that they focused on discrete relational structures whilst we constructed a continuous social space. Collectively, these evidence support the notion that grid-like codes underpin navigation in an abstract social cognitive map, be it discrete or continuous.

Intriguingly, our study found that the intensity of hexagonal modulation in precuneus aligned to estimated grid orientation of prefrontal region is negatively correlated with participants’ social avoidance tendency. While previous studies mainly reported grid-like codes in the entorhinal cortex and prefrontal cortex and its relevance to spatial navigation deficit in clinically risky population (Kunz et al., 2015), to the best of our knowledge, no report has shown the psychological relevance of grid-like code outside these classical regions to potential social deficit. One study of direct neuronal recording revealed grid-like neuronal activity in patients’ cingulate cortex during virtual spatial navigation task (Jacobs et al., 2013), implying that the human grid-cell network previously focused around entorhinal and prefrontal region also extends to the cingulate cortex. However, our study did not find evidence of grid-like pattern in precuneus. Thus, we cannot conclude that our result points to a direct relationship between precuneus grid-like code and social functionality. Meanwhile, precuneus, as a core node of the DMN, has abundant neuroanatomical connections with prefrontal regions (Greicius et al., 2009; Khalsa et al., 2014; Oane et al., 2020). Precuneus has also been reported to show a decrease in resting-state connectivity to parahippocampal gyrus and medial prefrontal cortex in patients with SAD (Yuan et al., 2018). Therefore, it is likely that downstream projections from the OFC/vmPFC motivated the observed correlation pattern in our study.

In conclusion, our present study demonstrates that navigating in a continuous social space recruited distance codes and grid-like codes in brain regions reported by spatial navigation studies and their behavioral and psychological relevance. Our findings further strengthen the notion that neural mechanisms involved in spatial cognitive map may play a domain-general role in maintaining abstract, structured representation of knowledge that support flexible cognitive behavior.

Our study has a few limitations. First, as we are using a visual analogue to guide participants to imaging moving in this abstract social space, grid-like coding could reflect processing of both the sensory information and the abstract social concept. It would be better to control for this as in Bao et al. (2019). Second, to make our study comparable to those investigating representation of abstract nonsocial knowledge, our scanner task investigate social navigation in a static environment and no social interaction or social decision is involved. Further studies should be conducted to investigate how such structured representation of social knowledge is reused and updated during social decisions and social interactions. Third, we specifically designed the abstract social value map to be formed by two orthogonal dimensions that have the same scale. This allows us to examine grid-like activity pattern with six-fold periodicity unambiguously. But this set-up may lack generality. For one thing, it is unlikely that all parts of the whole social value space span by the warmth and competence dimension are equally important. Social ties are more likely to be formed among human alike (McPherson et al., 2001) and human show the tendency to represent ingroup members more differentially than outgroup members (Hugenberg et al., 2010; Ostrom et al., 1993), so it is likely that some area of the social value space is represented in finer granularity than other. In spatial navigation, entorhinal grid cells showed sensitivity to geometric and environmental features, and the resulting distorted grid fields no longer form a regular hexagonal lattice and fail to show six-fold periodicity (Derdikman et al., 2009; He & Brown, 2019; Krupic et al., 2015). For instance, grid fields has been found to sometimes warp towards reward (Boccara et al., 2019). In this sense, spontaneous social value space may bear more resemblance to such kind of reward map covered by distorted grid fields rather than a regular, uniform square/circular open arena where six-fold symmetric grid-like coding was originally identified. For another, the geometric properties of social and non-social cognitive space in real life can be very different from that of the physical space in navigation. Our knowledge of other people in real life is often high-dimensional integrating multi-modal information from sensory features (face, voice etc.) to social cognitive information (hobbies, social status, etc.). Moreover, while dimensions of physical space (the cartesian axes) are orthogonal and their metrics are on the same scale, the dimensions of social perception in real life may not be orthogonal and may be incomparable. The stereotype content model predicted that there is a weak but low correlation between warmth and competence (Cuddy et al., 2009) and other studies have found there exists compensation between these dimensions (Kervyn et al., 2010). To the best of our knowledge, it remains elusive whether and how grid cells can represent such high-dimensional space with non-orthonormal basis and how the univariate and multivariate analysis pipeline used to identify grid-like coding in fMRI signal could be revised to address these concerns. Taken together, the intrinsic social cognitive map in which human structure their knowledge of others may be non-uniform, high-dimensional and have non-orthonormal dimensions. Future studies would need to take this into account and explore beyond the six-fold grid-like activity pattern in regular 2D space.

## Materials and Methods

### Participants

No power analysis was done to pre-determine the sample size. Instead, we tried to achieve a sample size comparable to previous studies investigating grid-like representations using fMRI (Doeller et al., 2010). 44 participants (18 males, mean age ± standard deviation: 21.59±2.56) recruited from surrounding universities finished all behavioral training and fMRI scanning in the current study at the Beijing Normal University Imaging Center for Brain Research (BICBR). Each participant underwent intensive behavioral training (including one learning session and one review session) and one fMRI scanning session **(Figure S1F)**. Three participants were excluded from all further analyses due to excessive head motion in the scanner (framewise displacement greater than 3mm). Another three participants failing to achieve satisfactory behavioral accuracies (lower than 60%) in the scanning session were excluded from all further analyses. The remaining 38 participants’ data were included in the behavioral and fMRI analysis reported in the main text (15 males, mean age ± standard deviation: 21.47±2.64).

The study was approved by the ethics committee of National Key Laboratory of Cognitive Neuroscience and Learning at Beijing Normal University (ICBIR_A_0071_011). All regulations were followed, and participants signed paper-form informed consent before the experiment. Participants received monetary compensation for their participation in the current study.

### Data acquisition

#### Behavioral data acquisition

Behavioral tasks were programmed using Psychophysics Toolbox-3 in MATLAB or E-prime 2.0. Online pilot rating study, pre- and post-experiment questionnaires were collected via Qualtrics. See **Supplementary Methods 1** for details on design and experiment procedure.

#### MRI data acquisition

We acquired T2-weighted functional images on a 3T SIEMENS MAGNETOM Prisma scanner with a 64-channel head coil. We acquired 33 slices, 3mm thick with: repetition time (TR) = 2000ms, echo time (TE) = 30ms, flip angle = 90°, field of view (FoV) = 224 mm, voxel size = 3.5×3.5×3.5 mm^3^. To correct for spatial distortion, a field map was acquired with dual echo-time images covering the whole brain with the following parameters: TR = 400ms, TE1= 4.92ms, TE2 = 7.38ms, flip angle = 60°, FoV = 224mm, voxel size = 3.5×3.5×3.5 mm^3^. A T1-weigted structural image was acquired with the following parameters: TR = 2530ms, TE= 2.98ms, flip angle = 7°, FoV = 256mm, voxel size = 0.5×0.5×1 mm^3^.

### Data analysis

All analyses were performed with customed code in MATLAB and R. Below is a brief summary of the analyses, see Supplementary Methods for details.

#### Behavioral Data analysis

Different performance indices were calculated for each task to test if performance improved over sessions (**Supplementary Methods 2.1**). We built linear mixed-effect models with random intercept and entered session as fixed effect otherwise specified.

#### Pre-processing of functional MRI data

Pre-processing of functional MRI data was done with SPM12. Functional images were spatially realigned to the first image in the time series and corrected for slice-timing. Spatial distortion based on field map. The T1-weighted structure image was co-registered to the mean aligned functional image, segmented and normalized to MNI space. The derived transformation parameters from structural image normalization were applied to normalize realigned functional images. Finally, smoothing was done with a 6mm full-width half-maximum Gaussian kernel.

#### Searching for representation of distance on 2D social value map in human fMRI signal

To look for neural representation of the travelled distance on the social value map, we conducted a whole-brain univariate analysis (**Supplementary Methods 2.2.3**). Two regressors were included in this GLM, one modeling the morph stage and one modeling the choice stage. The travelled Euclidian distance during morph stage was entered as a parametric modulator for the morph-stage regressor.

#### Searching for grid-like representations of 2D social value map in human fMRI signal

To search for evidence of entorhinal and prefrontal grid-like activity supporting representation of social space, we implemented two popular approaches to examine hexa-directional coding with fMRI data at the morphing stage in each trial.

The first one is the orientation-estimation approach (Bellmund et al., 2018; Doeller et al., 2010) based on univariate analysis (**Supplementary Methods 2.2.1-2.2.2**). In this approach, we first defined Regions of Interest (ROIs) that were sensitive to hexagonal modulation using univariate analysis. Thereafter, with a leave-one-out cross-validation procedure, we tested for hexagonal modulation aligned to grid orientation of the ROIs for each participant. In this procedure, we split the four acquired functional scanning runs into an ‘estimating set’ and a ‘testing set’. The estimating set consisted of three runs and was used to calculate the grid orientation of the ROIs. The remaining one run then served as the testing set where the alignment effect was tested with the inferred grid orientation. This procedure was repeated four times to independently yield the inferred orientations for each run. Specifically, trials were divided into 12 bins of 30° according to the offset of its trajectory direction from the putative grid orientation (**Figure 3A** left panel). The theoretical prediction for grid-like code is that neural signal should be stronger in trials aligned than misaligned to the grid orientation (0° modulo 60° vs 30° modulo 60°). In other words, activation should be higher in the aligned (odd-numbered) bins than the misaligned (even-numbered) bins (**Figure 3B** right panel). In addition to the above region-wise orientations, we also derived voxel-wise putative grid orientations in each ROI and used the circulate statistics to test whether there were clustered distributions of grid orientations as observed by the direct neuron recordings.

As we failed to find evidence of grid-like activity in EC aligned to its own putative grid orientation using the first approach, we took the anatomical mask of EC (Maass et al., 2015) and explored grid-like activity using a multivariate approach. This approach leverages representational similarity analysis (RSA) and is widely adopted based on the assumption that a distributed coding scheme is employed by the entorhinal cortex. Details of RSA analysis can be found in **Supplementary Methods 2.3**.

### Data Availability

We have published preprocessed smoothed and unsmoothed functional MRI data, the code to run the experiment and the code to run the univariate analysis reported in the main text. Dataset and code are publicly available at https://doi.org/10.57760/sciencedb.08637.

## Acknowledgements

We are grateful for the volunteers who participated in this study. We thank Alexandra Constantinescu for providing helpful information regarding the experiment design in their previous study. We thank Kelou Jin for his hard work and help when conducting pilot experiments. We thank Shen Zhang and Huagen Wang for their help in data analysis. This work was supported by the National Natural Science Foundation of China (32271092 and 32130045), the Major Project of National Social Science Foundation (19ZDA363), the Beijing Municipal Science and Technology Commission (Z151100003915122), the National Program for Support of Top-notch Young Professionals.

## Author contributions

Zilu Liang, Conceptualization, Investigation, Formal analysis, Visualization, Writing – original draft, Writing – review and editing; Simeng Wu, Investigation; Jie Wu, Investigation; Wenxu Wang, Conceptualization; Shaozheng Qin, Conceptualization, Supervision, Writing – review and editing; Chao Liu, Conceptualization, Supervision, Writing – review and editing, Funding acquisition.

## Notes

https://doi.org/10.57760/sciencedb.08637

### Competing Interest Statement

The authors have declared no competing interest.

### Summary of Updates

Rewrite parts of the method section to be more clear according to the reviewers' comments.

