## Supplementary Materials for "Social navigation: distance and grid-like codes support navigation of abstract social space in human brain"

### Supplementary Methods

#### 1. Design and experiment procedure

To look for neural underpinnings of navigation in an abstract social space, we adapted a set of tasks from previous studies illustrating the relevance of grid-like code in navigating abstract concept space . Participants received intensive training on navigating in this abstract social space with precision. They completed a learning session, a review session during behavioral training on the first day, and a scanning session on the second day **(Figure S1F)**.

*1.1 The social value map and experiment stimuli*

We designed the social value map, an abstract social space structuring one's social perception of other people's social value. It was defined by two ecologically important dimensions: competence and trustworthiness. A visualization with two adjacent bars enclosed by a square box was designed to represent the location on this social value map, with the height of each bar representing value of each dimension. We operationally defined these two dimensions under the framework of an investment game, in which trustee's social value is determined by the ability to earn profit (i.e., profit rate as competence) and the proportion of earnings returned to the investor (i.e., return rate as trustworthiness). Participants played the role of an investor in this investment task (**Figure S1A**) at the very beginning of the experiment to develop a concrete understanding of the visual analogue and the quantitative meaning of these two dimensions. Specifically, we limit the range of competence and trustworthiness between 0 and 1 to make it realistic, i.e., it is unlikely that a trustee will return you more than he/she has gained.

Six avatars, in analogy to landmarks in the physical environment, are placed on the social value map of size 1 unit × 1unit which is in accordance with the range of competence and trustworthiness. The current spatial arrangement aims to make avatars distinguishable on both dimensions while spreading widely across the whole space. Specifically, each participant's set of avatars' coordinates was sampled within circles of 1/30-unit radius around center coordinates (**Figure 1B**). Avatars were represented by standard photographs of volunteers who consented to the use of their photographs in the experiment.

To make sure the photographs did not induce prior perception of competence and trustworthiness, we conducted a pilot rating experiment online via Qualtrics in an independent sample of 39 participants (17 males, age: 22.56±2.41). On each trial, participants were shown one face and were asked to indicate their rating of the face on a 9-point Likert scale. Rating items were selected from the original SCM paper to reflect participants' perception of each faces' competence, trustworthiness, and attractiveness based solely on first impressions. Items reflecting competence included: ability, efficacy, and creativity. Items reflecting trustworthiness included: morality, friendliness, trustworthiness, and sincerity. Attractiveness was also included as an additional rating item. The order of rating items was randomized across trials to avoid habitual responses. We selected six (three from each gender) out of forty-eight faces that received middling ratings on all three dimensions (competence, trustworthiness, plus attractiveness) to serve as the stimuli in the current study. One sample t-tests revealed that ratings on all three dimensions of these six faces were not significantly different from five (the midpoint of the rating scale).

*1.2 Investment task*

At the beginning of the experiment, participants completed an investment task to develop a concrete understanding of the quantitative meaning of the two dimensions of the social value map (**Figure S1A**). The task is modified from the classic trust game. The investor is endowed with 1000 points and must allocate them between two agents. Competence is operationalized as how much more an agent can multiply his/her received investment while the return rate signals trustworthiness. The expectation of profit (the number of points) an investor can earn from one agent is thus formulated by:

$$E[Profit] = investment \times (1 + competence) \times trustworthiness \quad (1)$$

Participants were all assigned the role of investor. The two agents' competence and trustworthiness levels were preset by the experiment program and remained the same across participants.

*1.3 Match task*

Participants completed 30 trials of the match task to learn to use a non-spatial controller to morph the visualization (**Figure S1B**). One box with two bars appeared on the left side of the screen, and the participants were asked to use the controller to morph it to match the target box on the right side of the screen. The two boxes varied in the height of their enclosed bars, i.e., the competence and trustworthiness dimensions. The controller consisted of two horizontal thick black chunks, which signified how much the respective dimension would change. The closer to the top boundary, the more that dimension would change and vice versa. If placed on the midline, then the corresponding dimension would not change. Together, the controller's two chunks represented the ratio between how much the heights of two bars (i.e., values on two dimensions) changed relative to each other. To encourage participants to integrate two dimensions simultaneously, we instructed them to morph the bars by making as few transitions as possible and as accurate as possible. The morphing was continuous, with bars in the left box shrinking or stretching in height..

*1.4 Explore task*

Participants explored the vast social space by morphing the bars with different competence: trustworthiness ratio using the non-spatial controller to look for the six avatars (**Figure S1C**). No prior information about the characteristics of the avatars was provided to the participants. Thus, participants could only explore social space as if they were looking for landmarks in a newly introduced physical environment. An avatar would pop out when the visualization matched his/her characteristics. Specifically, the avatar would pop out when participants' current location fell within a 0.01-unit radius of the correct location on the social value map. In this way, participants not only learned the characteristics of each avatar but also became familiar with the whole space even though this map-like structure was never revealed to them.

*1.5 Collect Task*

After participants were fully acquainted with the avatars and the social space, they completed the collect task (**Figure S1D**). The task resembles the match task, except that instead of a target box, a target avatar appeared on the right side of the screen. Participants were instructed to morph the box on the left to match the characteristic of the avatar. Again, we instructed them to morph the bars by making as few transitions as possible and as accurate as possible. Each avatar was tested five times in each block, yielding 30 trials. Trials were presented randomly. Participants completed one block of collect task in each session.

#### *1.6 Recall task*

Participants completed the recall task both outside and inside the scanner (**Figure 1D**). On each trial of the recall task, participants were first shown a visualization morphing according to a predefined competence: trustworthiness ratio for 1 second. The bars then stopped morphing, and participants were instructed to imagine the bars keep morphing according to the same ratio, at the same speed, and for the same amount of time. After this, participants had to choose which of the three given options matched the bars after imagination.

Each block has 80 trials, each defined by a trajectory. To make sure trajectory directions were sampled uniformly on $[0,2\pi)$, we divided the whole range into 80 bins and each trial sampled a direction from one bin. Half of the trajectories led to learned avatars while the other half led to locations not associated with any avatar in the abstract space. Outside the scanner, participants completed two blocks of recall task in each session, resulting in 160 trials in total. In the scanner, participants completed four blocks of recall task, one in each run, resulting in 320 trials in total.

#### *1.7 Rank task*

We asked participants to rank the six learned avatars based on competence, trustworthiness, and willingness to cooperate with the avatar. When asking participants to rank based on willingness to cooperate, we specifically asked them to consider the two dimensions simultaneously and with the same weight.

#### *1.8 Compare task*

The compare task was designed to test whether participants formed an internal representation of the social value map, even though the map-like structure was never directly revealed to them during the experiment (**Figure S1E**). Participants were asked to compare two avatars on a given dimension and indicated which face had a higher value on the respective dimension using key press. The hypothesis was that if internal representation were formed, then it would take participants longer time to compare avatars with closer distance on the given dimension than to compare avatars far apart. Each possible combination of avatars was tested four times on each of the two dimensions. This yields 120 trials presented randomly. After that, an additional block on willingness to cooperate with 60 trials (four repetitions per combination) followed. Again, we specifically asked participants to give the two dimensions equal consideration when deciding their willingness to cooperate.

#### *1.9 Pre- and post-experiment face rating*

To reassure that the used face stimuli did not elicit biased social perception in the current sample, we asked our participants to rate the six avatars on a 9-point Likert scale when they signed up for the experiment (at least one day before the experiment). The rating items were identical to those in the online rating pilot experiment. To test whether participants learned each avatar's characteristics and updated their perception of the avatars, we asked our participants to rate the six avatars again after the fMRI scan.

*1.10 Map task*

At the end of the post-experiment questionnaire, participants were informed about the map-like structure and were asked to indicate the location of each avatar on an empty social value map using mouse click. We instructed them that the locations of avatars were defined by his/her level of competence and trustworthiness. We also asked participants whether they were aware of such a map-like structure and whether their strategy resembled this map-like organization.

**2. Data Analysis**

*2.1 Behavioral data analysis*

We calculated performance indices for different task. To test if performance improved over sessions, we built linear mixed-effect models with random intercept and entered session as fixed effect otherwise specified.

2.1.1 Explore task

First, we computed the time participants spent exploring the social space to find all six avatars. We predicted that participants would spend much less time in the review session than the learning session had they been fully acquainted with the space and the avatars. Next, the social space was divided into 15×15 subregions. We computed the amount of time spent in each subregion and plotted the corresponding color-coded trajectory maps. The top/bottom rows and leftmost/rightmost columns of subregions were classified as edges, yielding the index of "time at edges". In addition, "time at avatars" was computed based on the time the avatars were on screen. Note that all these timing indices were computed as a percentage of the total time spent navigating in the explore task in a given session. If participants were well-acquainted with the space and avatars, "time at edges" would decrease, and "time at avatars" would increase in the review session compared to the learning session. We built linear mixed-effect models to examine these hypotheses.

2.1.2 Collect task

First, we computed the mean number of transitions participants needed to morph the visualization to match the target avatar. Next, we computed the angle difference between participants' first transition and the ideal trajectory in each trial as the deviation from ideal trajectory. We further computed the mean deviation and the percentage of trials where participants deviated less than 15°. Last, we computed the mean distance between the target avatar and ending location indicated by the height of two bars in the visualization after participants finish morphing. Greater memory performance should be signaled by a lower mean number of transitions, a higher percentage of trials with deviation less than 15°, and a shorter mean distance from the target avatar.

2.1.3 Recall task

We computed the performance in the recall task in each session as the percentage of correct responses.

2.1.4 Compare task

We focused on the accuracy and response time in the compare task. We defined task-relevant distance as the distance between avatar pairs on the compared dimension. Specifically, for the two social value map dimensions, this was the distance on the competence/trustworthiness axis. For the willingness to cooperate block, this was the difference between the expected profit calculated in formula 1. We concatenated data from the review session and the scanning session and build separate linear mixed-effect models with random intercept for accuracy and response time. Session and task-relevant distance between avatar pairs was entered as fixed effect. We also included a random slope term for the task-relevant distance. The random slopes were extracted from the mixed effect model as indices for distance effect when exploring the behavioral relevance of spatial codes for the social value map.

2.1.5 Pre- and post-experiment face rating

We built three separate linear mixed-effect models with random intercept to test whether participants' ratings on competence, trustworthiness and attractiveness became more aligned with the avatars social characteristic after experiment. Three regressors were included as fixed effect: 1) time, i.e., Post- vs Pre-experiment; 2) Avatar's social characteristic; 3) interaction between time and avatar. Specifically, avatar's social characteristic was defined in alignment with the rating item entered as dependent variable. That is, in the competence rating regression model, social characteristic referred to the location on competence axis. Similarly, in the trustworthiness rating regression model, social characteristic referred to the location on trustworthiness axis. Finally, in the attractiveness rating regression model, expected profit was entered as social characteristic.

*2.2 Whole-brain univariate analysis*

All whole-brain univariate analyses were conducted in SPM12 following routine procedure. Particularly, we switched off SPM12's implicit threshold during model estimation and used an explicit mask from SPM12's default repository (the "mask_ICV.nii" file) that included all intracranial volumes instead. This specific treatment was done as we observed susceptible signal loss in the frontal and entorhinal region after applying the implicit threshold. Following routines from previous literature, the explicit mask was used instead (Ruge et al., 2019). In all univariate analyses, boxcar functions were used and the boxcar duration correspond to the duration of the modelled stage.

2.2.1 A functional localizer for hexagonal modulation: GLM1

GLM1 consisted of two regressors, one modeling the morph stage and one modeling the choice stage. The aim of GLM1 was to identify regions sensitive to hexagonal modulation. Let $\varphi$ denotes the hypothetical grid orientation of a region, and $\theta$ the trajectory (moving direction). If neural activity in a region is hexagonally modulated, then its activity should be a waveform of $\cos[6 * (\theta - \varphi)]$. Omitting error and intercept, this hypothesis can be expressed using the following formula:

$$\text{Activity} = \omega * \cos[6 * (\theta - \varphi)]$$

$$= \omega * cos6\varphi cos6\theta + \omega * sin6\varphi sin6\theta \quad (2)$$

$$= \beta_{cos6\theta} cos6\theta + \beta_{sin6\theta} sin6\theta \quad (3)$$

where:

$\beta_{cos6\theta} = \omega * cos6\varphi$

$\beta_{sin6\theta} = \omega * sin6\varphi$

If the theoretical prediction that there is an effect of hexagonal modulation is true, $\omega$ should be significantly different from zero.

Based on this, a pair of sine and cosine regressors were used as parametric modulators for the morph stage. In this way, testing against the null hypothesis of $\beta_{cos6\theta} = \beta_{sin6\theta} = 0$ is essentially testing against the null hypothesis of $\omega = 0$. Therefore, at individual-level, F-test was used to search for potential regions modulated by a linear combination of $\beta_{cos6\theta} cos6\theta + \beta_{sin6\theta} sin6\theta$. These individual F-statistics were transformed into Z-statistics before entering group-level one-sample t-test. In future correlation analyses, the Z-statistics were extracted as an index for hexagonal modulation effect.

2.2.2 Iterative cross-validation analysis for hexagonal consistency: GLM2

By binning trials according to trajectory's alignment to a putative grid orientation phi, GLM2 aimed to test for hexagonal consistency.

In this cross-validation procedure, we separated each participants data into two set, an estimating set with three runs and a testing set with one run. The estimating set was used to calculate the grid orientation of the ROIs. The remaining one run then served as the testing set where the alignment effect was tested with the inferred grid orientation. The theoretical prediction for hexagonal consist grid-like code is that neural signal should be stronger in trials aligned than misaligned to the grid orientation.

To estimate grid orientation, GLM1 was applied to the estimating set yielding beta estimates for the sine and cosine regressors ($\beta_{sin6\theta}$ and $\beta_{cos6\theta}$) in each voxel. We extracted these beta estimates from ROI masks. Then, within a given ROI, beta estimates averaged across all voxels within this region for sine and cosine regressors respectively. The averaged beta estimates ($\overline{\beta_{sin}}$ and $\overline{\beta_{cos}}$) were used to calculate the grid orientation for this region (grid orientation $\varphi = \left[artan(\frac{\overline{\beta_{sin}}}{\overline{\beta_{cos}}})\right]/6$).

To test for the prediction of consistency effect, we classified trials according to $\theta$'s (trajectory direction) offset from $\varphi$ into 12 bins of 30° (**Figure 3A left panel** in the main text) , yielding 6 bins of aligned trials (0° modulo 60°) and 6 bins of misaligned trials (30° modulo 60°).

The above steps yielded the grid orientation applied to classify trials in the testing set. Each run acted as estimating set three times and received putative grid orientation as testing set once. Eventually, the four testing sets were fitted with GLM2. GLM2 consisted of 12 regressors, each modelling the morph stage of one bin of trials. At individual level, contrast was built to test the activity difference between aligned and misaligned trials (align>misalign). These first-level contrasts were entered into second level one-sample t-test to look for to regions that show hexagonal consistency.

2.2.3 Identifying distance code with parametric modulation: GLM3

GLM3 resembled GLM1, which also consisted of two regressors, one modeling the morph stage and one modeling the choice stage. The aim of GLM3 was to identify regions sensitive to distance modulation. Thus, we entered the travelled Euclidian distance during morph stage as a parametric modulator for the morph-stage regressor. At individual-level, t-contrast was built to search for potential regions modulated by the distance parametric modulator. These contrasts were entered into second level one-sample t-test to look for to regions that represent distance during social navigation.

2.2.4 Behavioral relevance of spatial codes

To specifically test the effect of grid-like code in regions that survive correction in GLM2 on behavior, we extracted the beta estimates of align>misalign contrast for each region and for each participants. Then we computed Pearson correlation between these beta estimates and accuracy as well as response time in the recall task in scanner. This procedure was applied to the regions demonstrating distance representation (GLM3) as well. As multiple tests were performed, we adopted FDR correction to p-values to lower false discovery rate. No significant correlation was found (**Figure S7**).

For whole-brain analyses to explored if there is any correlation between neural indices of map-like representation and behavior performance outside the scanner as well as individual differences, we calculated two categories of behavioral indices to explore the relevance of the spatial codes calculated above. The first category of indices reflects participants ability to make decision based on the social value map. Specifically, we considered distance effect during the cooperation block when participants decide the avatar they are more willing to cooperate with when taking into account both dimensions. Distance effect indices were extracted from the linear mixed effect model estimates specified in the behavioral data analysis section. The second category of indices reflects participants' social trait, including social anxiety and social avoidance scores. We explored the behavioral relevance of grid-like code by entering the above indices as covariates into the second-level analysis when testing grid orientation consistency effect (GLM2), and all covariates were tested in separate regression models.

*2.3 Multivariate analysis in Regions of Interest*

As we failed to find evidence of grid-like activity in EC aligned to its own putative grid orientation using the first approach, we took the anatomical mask of EC (Maass et al., 2015) and explored grid-like activity using a multivariate approach. This approach is widely adopted based on the assumption that a distributed coding scheme is employed by the entorhinal cortex. We conducted conventional representational similarity analysis on the unsmoothed data as implemented by CosmoMVPA toolbox (Oosterhof et al., 2016) in MATLAB.

First, we estimated trial-specific activation for each trial in each run. As trajectories were drawn randomly in the social space, each trial was defined by a unique trajectory, yielding a rich-condition design. It has been suggested that in such fast event-related (fast ER) designs, signal for nearby trials tend to overlap in time. Conventional approach in block or slow ER design that includes each trial as a separate regressor in a large model may be problematic under fast ER settings as estimates can become unstable due to collinearity between the trial-specific regressors. To obtain more accurate estimate of trial-specific activation, we leveraged the 'Least Squares Separate' (LSS) approach (Abdulrahman & Henson, 2016; Mumford et al., 2012). This approach built separate GLMs for each trial with one target regressor modeling the current trial of interest and a second nuisance regressor modeling the rest. After that, we took the beta estimate of the target regressor the as the estimation of trial-specific activation. Signals were extracted from the entorhinal ROIs (from entorhinal as a whole or from four subregions) as trial-specific activity pattern for each of the 320 trials (**Figure S4A**).

We then tested whether this multivariate pattern is hexagonally modulated. Though the exact implementations varied slightly across studies, they fell into the following two categories according to their dependence on the estimated putative grid orientation.

The 'orientation-independent' approach (**Figure S4B**) tested the hypotheses that multivariate pattern similarity between trajectory pairs is proportional to the angular difference between their moving direction modulated by 60° (Bellmund et al., 2016; Vigano & Piazza, 2020; Vigano et al., 2021). We first calculated the representational similarity between all possible pairs of trials, yielding a 320×320 dissimilarity matrix (DSM) for neural data (**Figure S4A**). Then, we derived model DSM from theoretical prediction (**Figure S4B**). Finally, we computed Spearman's rank correlation between neural DSM and model DSM for each participant. One-sample t-test were conducted across participants on the Fisher z-transformed correlation coefficients to test whether the correlation was significantly above zero.

The 'orientation-dependent' approach **(Figure S4C)** leveraged the assumption that if activity in a given ROI is aligned to a preferred orientation, the similarity of multivoxel patterns in this region should be higher for aligned trial-pairs than that between aligned and misaligned trial-pairs (Bao et al., 2019). We computed the grid-orientation in each entorhinal sub region using the leave-one-out cross-validation procedure (**Supplementary Methods 2.2**). Trials were then classified into aligned and misaligned trials according to the angular difference between trajectories and the estimated grid orientation. We then computed the similarities between aligned-aligned trial pairs (AA) and aligned-misaligned trial pairs (AM). By subtracting similarity between AM pairs from similarity between AA pairs, we yield a pattern similarity difference index for each participant (**Figure S4C**). Paired t-tests were conducted to test whether the differences were significantly greater than zero. As multiple tests were performed, we adopted FDR correction to p-values to lower false discovery rate.

#### *2.4 Testing distribution of grid orientation*

All tests in this section are circular statistics *(Mardia & Jupp, 1999)* implemented by the MATLAB toolbox *CircStat (Berens, 2009)*.
To test if the grid orientations within an ROI is similar, we calculated the putative grid orientations for each voxel in an ROI using voxel-wise beta estimates of the sine and cosine regressors from GLM1. Then the vector of putative voxel-wise grid orientations is entered into a V-test to test if the distribution of the orientations is clustered. We specifically used the V-test instead of the Rayleigh's test because the alternative hypothesis in V-test assumed that data have a known mean direction. This is more in line with the hypothesis that neighboring grid cells share similar grid orientation. As the cross-validation procedure yields three estimating sets for each participant, this procedure is repeated for all estimating sets and for all participants.
To test the distribution of grid orientation in the population, for one ROI, we calculated the grid orientation for each participant using the averaged beta estimates of the sine and cosine regressors from GLM1 in this ROI. This vector of grid orientation is entered into a Rayleigh's test. Following previous study, we assume that the grid orientations across participants would be uniformly distributed.

*2.5 Relationship between temporal sigal-to-noise ratio (tSNR) and hexagonal modulation effect*

Following previous literature (Murphy et al., 2007; Triantafyllou et al., 2005), we calculated the voxel-wise tSNR of a timeseries, i.e. the tSNR for a single voxel in a single run of one participant, as:

$$tSNR = \frac{\mu}{\sigma}$$

$\mu$ is the mean activity of the timeseries, and $\sigma$ is its standard deviation.

The tSNR of each voxel is calculated as the mean across four runs, and the tSNR for each ROI is calculated as the averaged tSNR of all voxels in the ROI.

To test the relationship between tSNR and hexagonal modulation effect, we conducted correlation analysis across participants and within each participant. For the across-participant analysis, we tested the correlation between the mean hexagonal modulation effect in an ROI and its mean tSNR across all participants. For the within-participants analysis, for each participant, in an ROI, we calculated the correlation coefficient between voxel-wise hexagonal modulation effect and voxel-wise tSNR. Then, after Fisher Z transformation, we tested whether the participant-level correlation coefficients were significantly different from zero using Wilcoxon signed-rank test at group level.

### Supplementary Figures

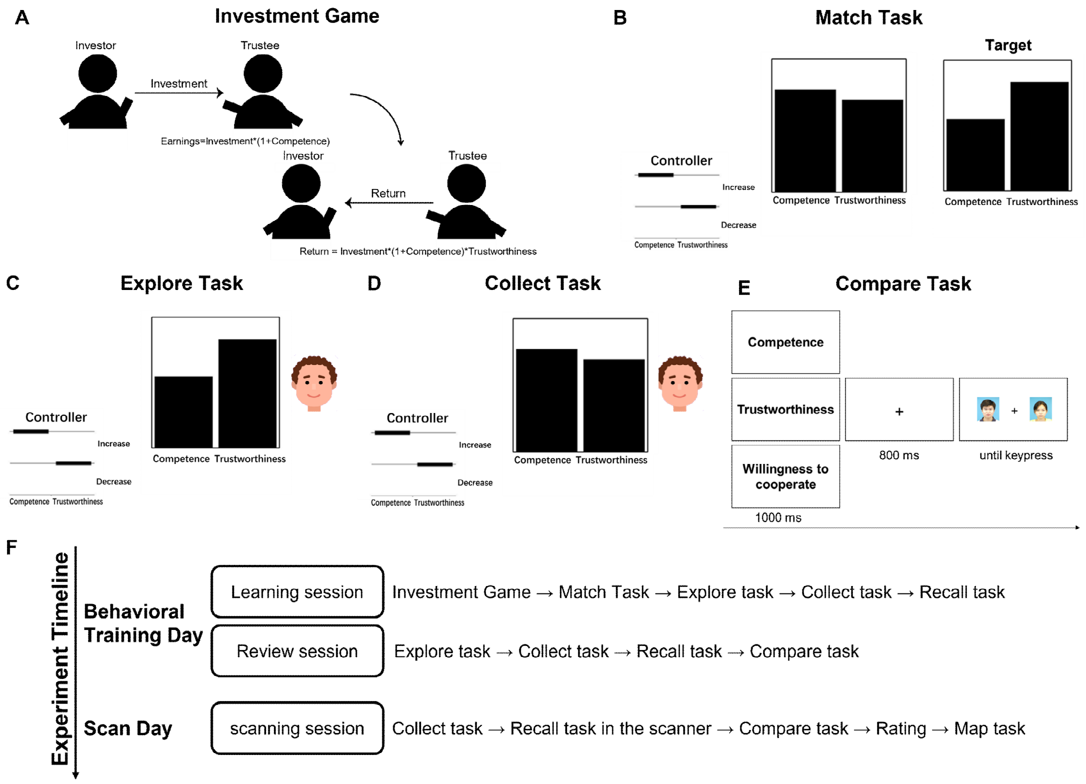


**Figure S1. Behavioral Training Tasks.** ***(A)*** *Schematic illustration of the investment game and its relation to the competence and trustworthiness dimensions.* ***(B-D)*** *Example screenshot of match, explore and collect task.* ***(E)*** *Timeline of compare task.* ***(F)*** *Timeline of the experiment and tasks completed by participants in each session.*

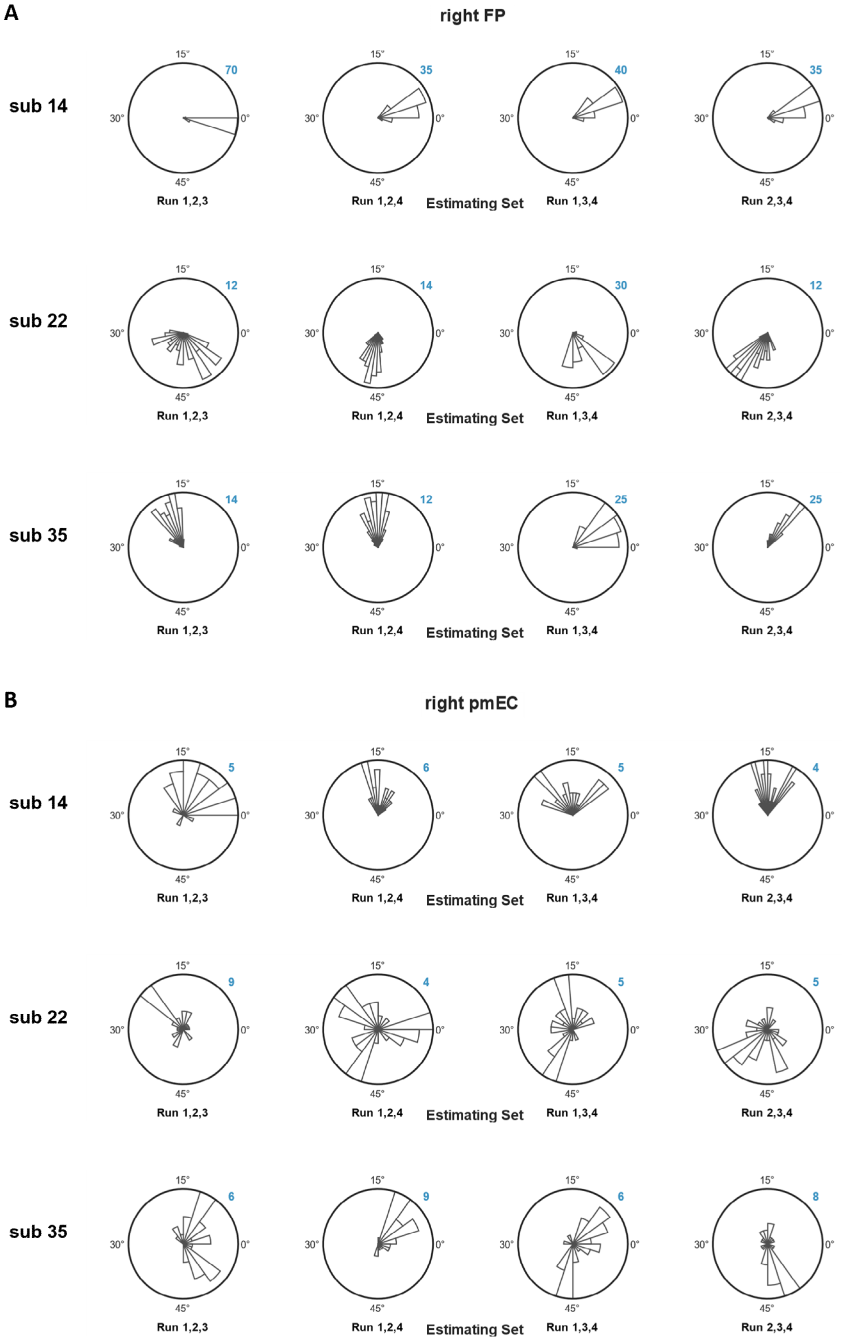


**Figure S2. Distribution of voxel-wise grid orientation of example participants** (voxel-wise distribution plot of all participants can be viewed at https://doi.org/10.57760/sciencedb.08637). *Blue number indicates voxel count in the bin with most voxels.*

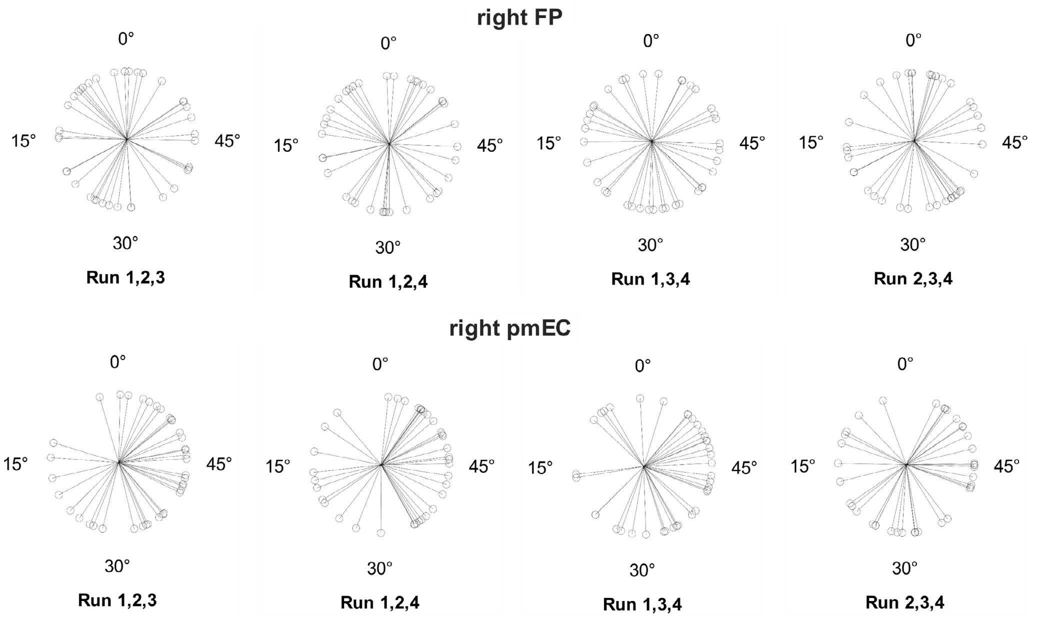


**Figure S3. Distribution of grid orientation across participants.** *Each point is the estimated grid orientation in a given estimating set in a given ROI for one participant.*

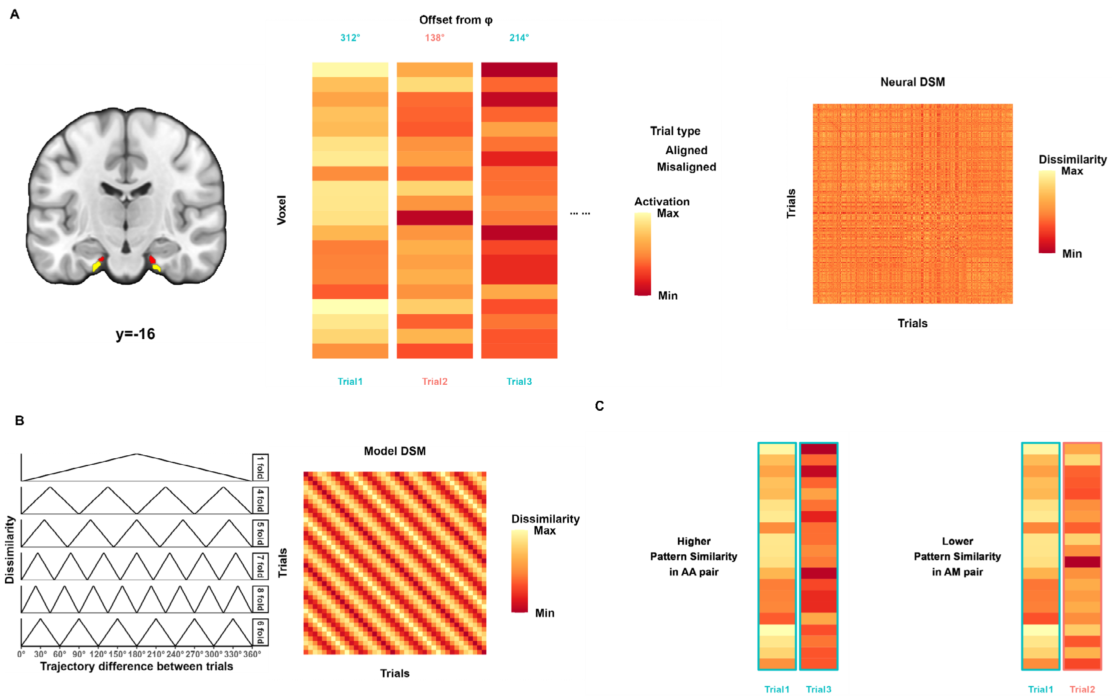


**Figure S4. Analysis pipeline of multivariate pattern analysis in entorhinal ROI.** *(A) Signals from four subregions of Entorhinal cortex were extracted. (**B**) Model-based analysis Hexagonal consistency effect in the entorhinal subregions. (**C**) Model-free analysis (angle independent) RSA results in the whole entorhinal cortex and different entorhinal subregions.*

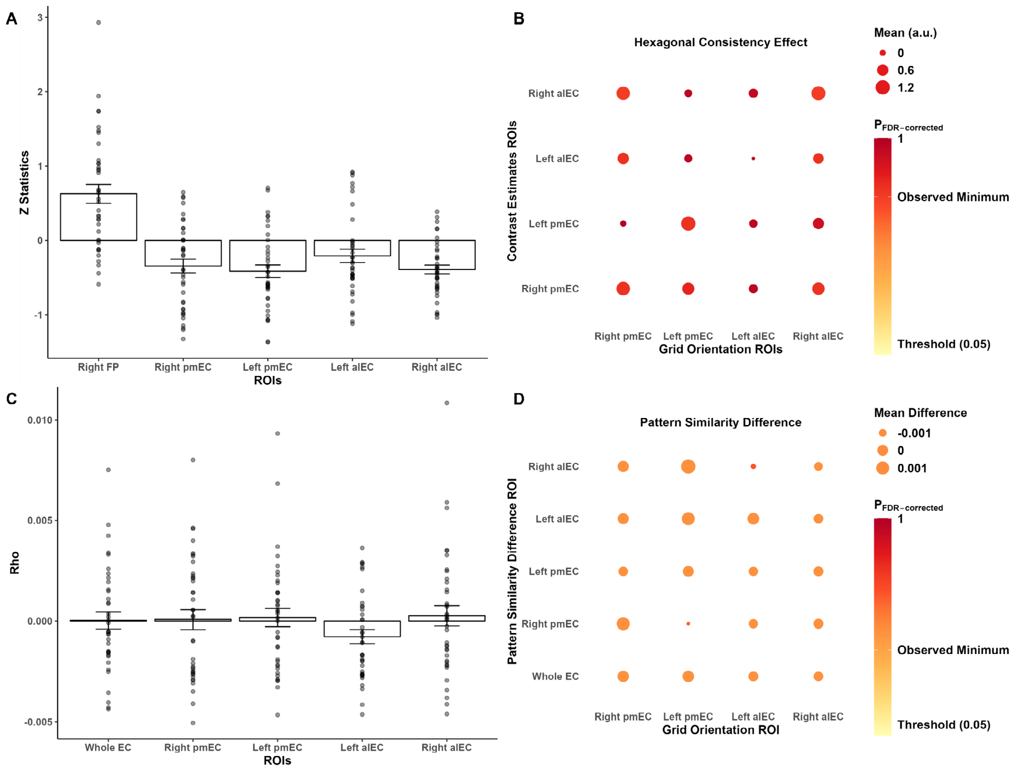


***Figure S5. ROI analysis of univariate and multivariate grid-like code in the entorhinal cortex.*** *(A) Z-transformed F statistics of hexagonal modulation effect in the entorhinal sub regions in comparison with the Frontal Pole ROI. (**B**) Hexagonal consistency effect in the entorhinal sub regions. (**C**) Model-based (*orientation-independent*) RSA results in the whole entorhinal cortex and different entorhinal sub regions. (**D**) Model-free (*orientation-dependent*) RSA results in the entorhinal sub regions. Pattern similarity difference between aligned and misaligned trials based on putative grid orientations from different entorhinal subregions. Abbreviations: FP, frontal pole; pmEC, posterior-medial entorhinal cortex; alEC, anterior-lateral entorhinal cortex.*

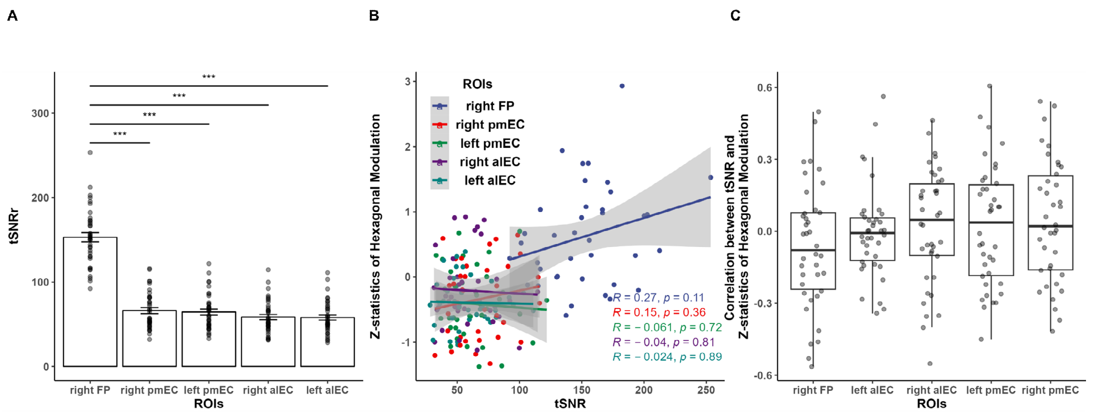


**Figure S6. Relationship between tSNR and the strength of evidence of hexagonal modulation effect in frontal pole and entorhinal ROIs.** *(**A**) Across participants, mean tSNR in right Frontal Pole ROI is higher than all four subregions of Entorhinal cortex. (**B**) Across participants, in each of the five ROI, its mean tSNR is not correlated with the Z-statistics of hexagonal modulation effect. (**C**) Within participants, in each of the five ROI, voxel-wise tSNR is not correlated with the Z-statistics of hexagonal modulation effect.*

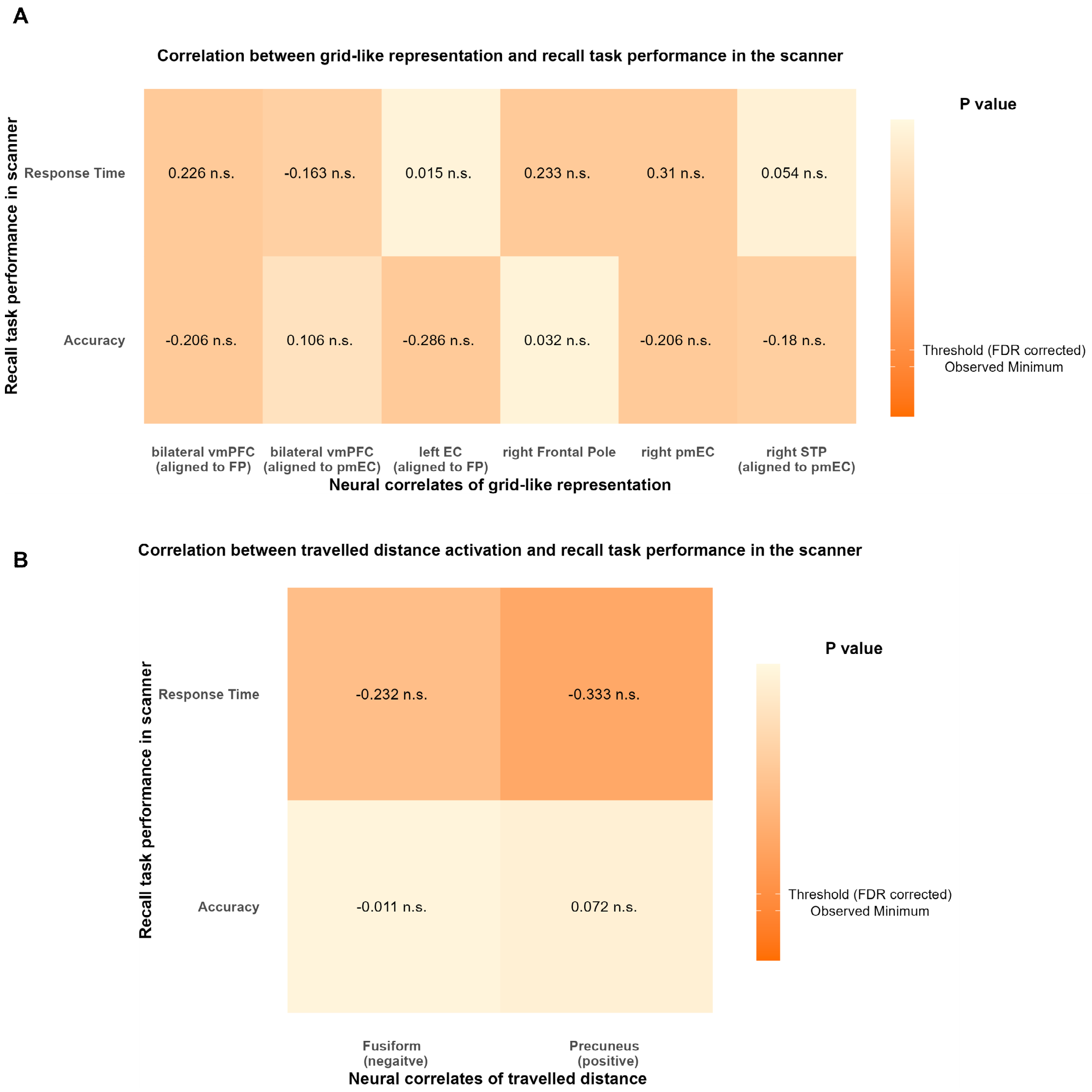


**Figure S7. No evidence of correlation between grid-like *(A)* and distance *(B)* representation and performance in the scanner.** *Number in each grid indicates Pearson correlation coefficient between a pair of behavioral and neural variable, color indicates p-value result from statistical test (FDR-corrected).*

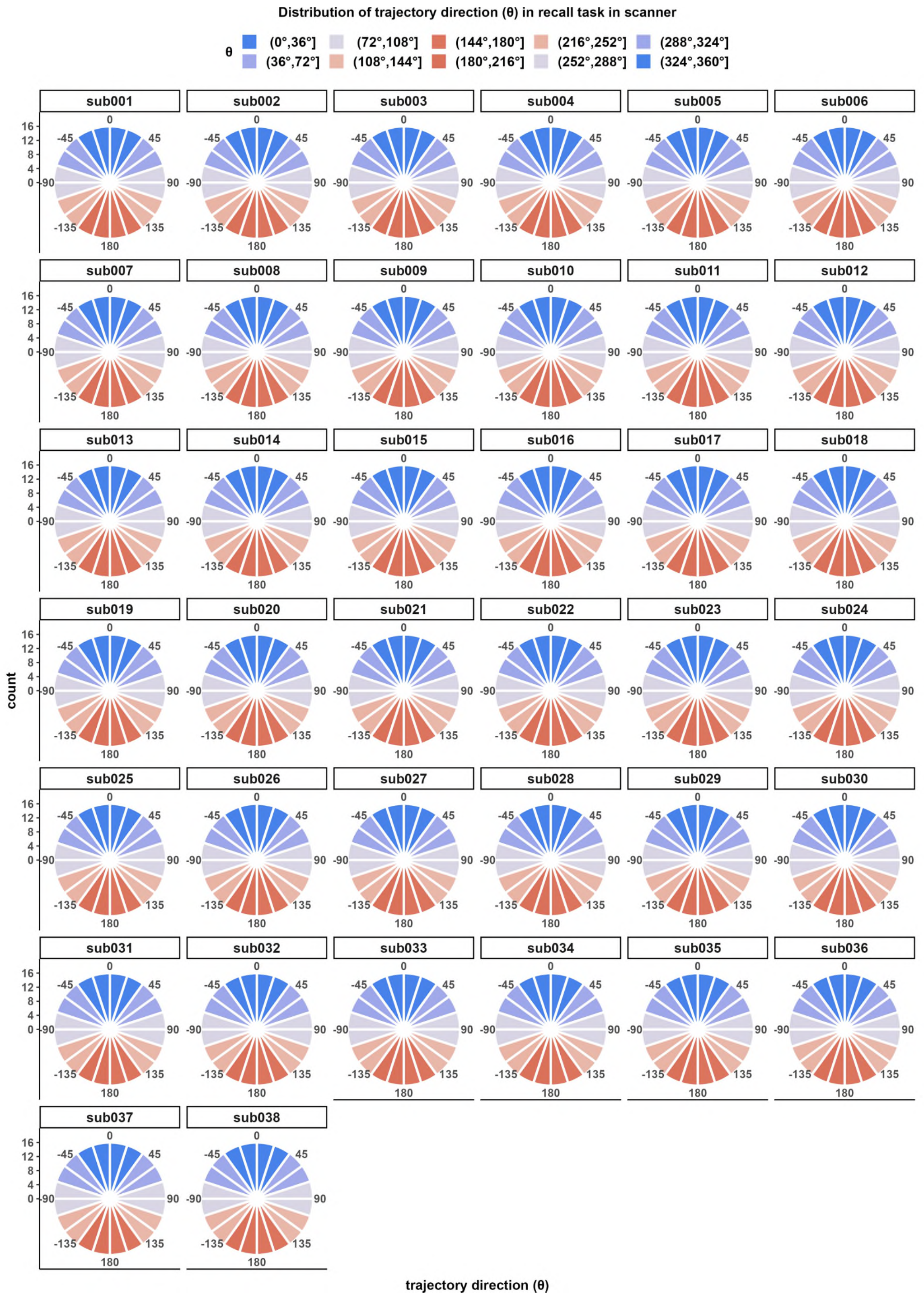


**Figure S8. Distribution of trajectory direction (theta) in the recall task in scanner.** *Trajectory directions from all four runs were divided into 20 bins to plot histogram. Angle of polar plot indicate trajectory direction, distance from center indicate number of trials in the bin.*

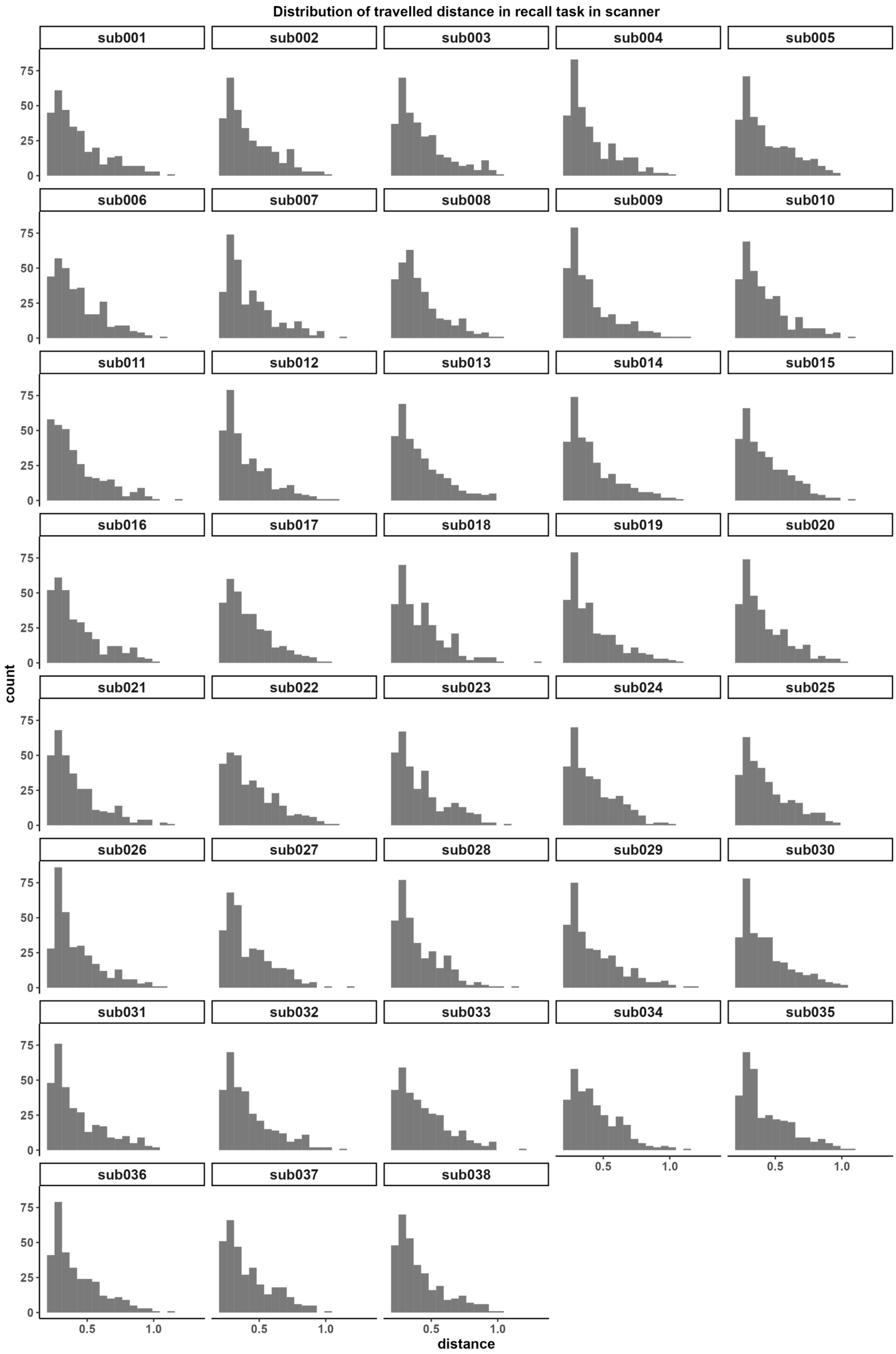


**Figure S9. Distribution of trajectory length (travelled distance) in the recall task in scanner.**

*Trajectory lengths from all four runs were divided into 20 bins to plot histogram.*

**Supplementary Tables**

**Table S1 Related to Figure 1H.**

**A. Linear Mixed Effect Model for different rating items**

| | Competence | Trustworthiness | Attractiveness |
|---|---|---|---|
| (Intercept) | 5.157 ***<br>[4.775, 5.539] | 5.614 ***<br>[5.276, 5.951] | 4.590 ***<br>[4.168, 5.011] |
| Post vs Pre | -2.173 ***<br>[-2.707, -1.639] | -1.998 ***<br>[-2.469, -1.527] | -0.828 **<br>[-1.364, -0.292] |
| Avatar | -0.395<br>[-1.024, 0.234] | -0.637 *<br>[-1.240, -0.034] | -0.281<br>[-0.710, 0.149] |
| avatar * (Post vs Pre) | 4.812 ***<br>[3.923, 5.702] | 5.185 ***<br>[4.333, 6.038] | 1.964 ***<br>[1.357, 2.572] |
| N (observation) | 456 | 456 | 456 |
| N (id) | 38 | 38 | 38 |
| AIC | 1600.834 | 1589.456 | 1808.221 |
| BIC | 1625.569 | 1614.191 | 1832.956 |
| R2 (fixed) | 0.302 | 0.337 | 0.129 |
| R2 (total) | 0.313 | 0.347 | 0.223 |

*** p < 0.001; ** p < 0.01; * p < 0.05.

**B. Follow-up analysis of interaction term in mixed effect models: Simple Slope of avatar in different sessions**

| Rating Item | Moderator levels<br>session | Estimate<br>[lower CI, upper CI] | SE | t (415) | p |
|---|---|---|---|---|---|
| Competence | Pre-experiment | -0.395<br>[-1.026, 0.236] | 0.321 | -1.231 | 0.890 |
| | Post-experiment | 4.417<br>[3.786, 5.048] | 0.321 | 13.766 | <0.001 |
| Trustworthiness | Pre-experiment | -0.637<br>[-1.241, 0.032] | 0.308 | -2.069 | 0.980 |
| | Post-experiment | 4.549<br>[3.944, 5.154] | 0.308 | 14.787 | <0.001 |
| Attractiveness | Pre-experiment | -0.281<br>[-0.711, 0.150] | 0.219 | -1.281 | 0.900 |
| | Post-experiment | 1.684<br>[1.253, 2.114] | 0.219 | 7.684 | <0.001 |

Statistic results for right-sided t test against zero (noninferiority)

**Table S2. Related to Figure 2. Neural codes representing travelled distance on the social value map**

**A. Regions positively correlated with travelled distance.**

| Anatomical Description | Hemisphere | Peak MNI coordinates x | y | z | Peak t value | Cluster Size | $p_{FWE}$ |
|---|---|---|---|---|---|---|---|
| Precuneus | R | 12 | -56 | 26 | 4.530 | 151 | 0.054 |
| Precuneus | L | -12 | -54 | 18 | 4.952 | 134 | 0.082 |

**B. Regions negatively correlated with travelled distance.**

| Anatomical Description | Hemisphere | Peak MNI coordinates x | y | z | Peak t value | Cluster Size | $p_{FWE}$ |
|---|---|---|---|---|---|---|---|
| Fusiform | R | 36 | -48 | 20 | 4.530 | 766 | <0.001 |
| Fusiform | L | -38 | -56 | -10 | 4.952 | 234 | 0.008 |
| Middle occipital gyrus | R | 42 | -80 | 8 | 4.952 | 196 | 0.018 |

**Table S3. Regions showing hexagonal modulation (GLM1, Supplementary Methods 2.2.1)**

| Anatomical Description | Hemisphere | Peak coordinates (MNI) | | | Peak t value | Cluster | |
|---|---|---|---|---|---|---|---|
| | | x | y | z | | Size | $p_{FWE}$ |
| Superior parietal gyrus | R | 28 | -66 | 56 | 5.021 | 522 | <0.001 |
| Precuneus | L | -6 | -46 | 38 | 5.707 | 426 | <0.001 |
| Middle frontal gyrus | R | 42 | 24 | 34 | 5.221 | 271 | <0.001 |
| Paracentral Lobule | R | 10 | -36 | 68 | 4.835 | 120 | <0.001 |
| Middle frontal gyrus | R | 38 | 42 | 30 | 4.849 | 103 | 0.001 |
| Frontal pole | L | -26 | 50 | 0 | 4.269 | 71 | 0.014 |
| Frontal pole | R | 26 | 52 | -2 | 4.494 | 69 | 0.017 |
| Angular | L | -52 | -60 | 26 | 4.718 | 58 | 0.041 |

**Table S4. Related to Figure S6. Group-level Wilcoxon signed-rank test of correlation between voxel-wise tSNR and the Z-statistics of hexagonal modulation effect in frontal pole and entorhinal ROIs.**

| ROIs | N | Test Statistics | p |
|---|---|---|---|
| right Frontal Pole | 38 | 268.000 | 0.140 |
| left anterior-lateral entorhinal cortex | 38 | 331.000 | 0.572 |
| right anterior-lateral entorhinal cortex | 38 | 415.000 | 0.528 |
| left posterior-medial entorhinal cortex | 38 | 396.500 | 0.712 |
| right posterior-medial entorhinal cortex | 38 | 433.000 | 0.369 |
